# EMMA: A New Method for Computing Multiple Sequence Alignments given a Constraint Subset Alignment

**DOI:** 10.1101/2023.06.12.544642

**Authors:** Chengze Shen, Baqiao Liu, Kelly P Williams, Tandy Warnow

## Abstract

**Background:** Adding sequences into an existing (possibly user-provided) alignment has multiple applications, including updating a large alignment with new data, adding sequences into a constraint alignment constructed using biological knowledge, or computing alignments in the presence of sequence length heterogeneity. Although this is a natural problem, only a few tools have been developed to use this information with high fidelity.

**Results:** We present EMMA (Extending Multiple alignments using MAFFT-add) for the problem of adding a set of unaligned sequences into a multiple sequence alignment (i.e., a constraint alignment). EMMA builds on MAFFT-add, which is also designed to add sequences into a given constraint alignment. EMMA improves on MAFFT--add methods by using a divide-and-conquer framework to scale its most accurate version, MAFFT-linsi--add, to constraint alignments with many sequences. We show that EMMA has an accuracy advantage over other techniques for adding sequences into alignments under many realistic conditions and can scale to large datasets with high accuracy (hundreds of thousands of sequences). EMMA is available at https://github.com/c5shen/EMMA.

**Conclusions:** EMMA is a new tool that provides high accuracy and scalability for adding sequences into an existing alignment.

## 1 Background

### 1.1 Adding Sequences to a Multiple Sequence Alignment

Multiple sequence alignment (MSA) is a crucial precursor to many downstream biological analyses, such as phylogeny estimation [1], RNA structure prediction [2], protein structure prediction [3], etc. Obtaining an accurate MSA can be challenging, especially when the dataset is large (i.e., more than 1000 sequences). In some cases, the problem of estimating an alignment on a large dataset can be addressed through approaches that seek to add sequences into a given alignment, without allowing the given alignment to change; for this reason, the given alignment is referred to as a “constraint alignment”. For example, biological knowledge can be used to form a reference alignment on a subset of the sequences, and then the remaining sequences can be added to the reference alignment; this has the potential to improve accuracy compared to methods that do not include biological knowledge to define a constraint alignment. Another case where adding sequences into an existing alignment occurs is when new sequences or genomes are added to databases, leading to the opportunity to add the new sequences for each gene in the genome into a growing alignment. In this second case, adding sequences into the existing alignment avoids the need to recompute the alignment from scratch and could lead to substantial running time benefits. A third case is for *de novo* multiple sequence alignment, where a subset of the sequences is selected and aligned, and then the remaining sequences are added into this “backbone alignment”; examples of such methods include UPP [4], UPP2 [5], WITCH [6], WITCH-ng [7], HMMerge [8], and MAFFT-sparsecore [9]. One of the motivations for this type of alignment method (which we refer to as “two-stage” methods) is when the input dataset has substantial sequence length heterogeneity, which can result in poor alignment accuracy using standard methods [4]. Thus, adding sequences into existing alignments is a natural problem with multiple applications to biological sequence analysis.

A few methods have been developed to add sequences into an existing alignment, with MAFFT--add [10] perhaps the most well-known. However, multiple sequence alignment methods that operate in two steps – i.e., they first extract and align the backbone sequences and then add the remaining sequences into this backbone alignment – can be modified to enable sequences to be added into a user-provided alignment.

### 1.2 HMM-based methods

Many of the methods that have been developed for adding sequences into a given multiple sequence alignment operate by representing the existing alignment by either a single HMM or by an ensemble of HMMs. Then, for every additional sequence, which we call “query sequences”, the HMM or ensemble of HMMs is used to find an alignment of the query sequence to the given alignment. Examples of such approaches include functionality provided in UPP [4], UPP2 [5], WITCH [6], WITCH-ng [7], and HMMerge [8]. We refer to these functions by appending “-add” to the MSA method name (e.g., this functionality in WITCH is referred to as WITCH-add). In this study, we examine the performance of WITCH-ng-add as it has been shown to be at least as accurate and generally faster than WITCH-add, and both are in turn at least as accurate as UPP-add and UPP2-add. Finally, HMMerge-add is slower than WITCHng-add and so is omitted from this study. See Supplementary Materials Section S1 for additional details about these HMM-based methods.

Because HMM-based methods operate by aligning the query sequences to one or more HMMs, homologies between query sequences can only be discovered if these homologies align through match states in the HMMs. This approach has the potential to miss valid homologies between query sequences, for example when the given alignment is insufficiently representative of the entire family. Thus, only query sequence characters (i.e., nucleotides or residues) that are aligned through match states in the HMMs can be placed in columns that have other nucleotides or residues; if the character is added to the alignment through an insertion state, it will never be detected as homologous to any other character in any other sequence. This aspect of using HMMs for alignment has a potentially significant impact on the accuracy of the alignments of query sequences to the backbone alignment.

### 1.3 MAFFT--add and MAFFT-linsi--add

MAFFT--add [10] in its default setting uses a standard progressive alignment procedure with two iterations to add query sequences. In each iteration, it computes the pairwise distance matrix between the complete set of sequences (both in the backbone and the query sequence set) using shared 6-mers. Then, it computes a guide tree using the distance matrix and builds an alignment. More specifically, for each node in the guide tree, MAFFT--add does an alignment computation only if a query sequence is involved at the node (i.e., at least one child has some query sequences). Otherwise, it simply uses the alignment from the backbone, and so is guaranteed to preserve the input backbone alignment. This property of guaranteeing that the input backbone alignment is not changed is true of the more accurate variants of MAFFT--add, including the variant that uses MAFFT-linsi to add query sequences (which we refer to as MAFFT-linsi--add), which we briefly describe below.

MAFFT-linsi--add has two differences to the default version of MAFFT--add. First, MAFFT-linsi--add uses *localpair* (local pairwise alignment scores) for the distance matrix calculation, which is more accurate than shared 6-mers. Second, it only runs for one iteration of progressive alignment and uses at most 1000 iterations of iterative refinement after the progressive alignment finishes. MAFFT--add and MAFFT-linsi-add have runtimes that are at least quadratic in the input size due to the *O*((*m* + *q*)^2^) distance matrix calculation, where there are *q* query sequences and *m* sequences in the provided backbone alignment. MAFFT-linsi--add is even less scalable since its distance calculation is more costly. In addition, MAFFT-linsi--add does many steps of refinement that further impact the runtime. Hence, the developers of MAFFT recommend that MAFFT-linsi--add be limited to a few hundred sequences [12].

**Figure 1:**
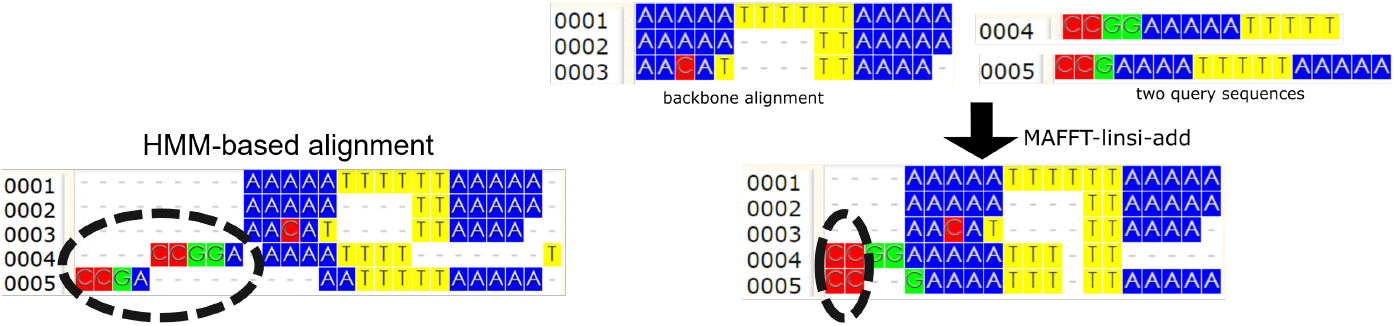
Comparing MAFFT-linsi--add to UPP-add. We ran MAFFT-linsi--add and UPP-add on a backbone alignment and two query sequences. The initial two sites in the MAFFT-linsi--add alignment each contain two letters from the query sequences, thus indicating it detects them as homologous to each other, although there are no letters from the backbone alignment in these two sites. This is impossible for any method that represents the backbone alignment by one or more HMMs, and then adds sequences to the backbone alignment using the HMMs. Thus, UPP-add places these initial letters of the query sequences into separate sites. Visualization is done with WASABI [11].

### 1.4 Limitations of HMM-based methods

Both MAFFT-linsi--add and MAFFT--add can recover homologies between query sequences that do not have homologous characters in the backbone alignment, and – as mentioned above – they are guaranteed to leave the backbone alignment unchanged as they add query sequences.

In contrast, methods such as WITCH-add, WITCH-ng-add, etc., that use HMMs or ensembles of HMMs to represent the backbone alignment cannot find homologies between letters in the query sequences that do not also have homologs in the backbone alignment. This is an inherent limitation of HMM-based methods for adding sequences into backbone alignment and implies that MAFFT--add and MAFFT-linsi--add may be more robust to the selection of the backbone sequences than the HMM-based methods.

Figure 1 gives an example of a backbone alignment and query sequences, comparing MAFFT-linsi--add and UPP-add. Note that MAFFT-linsi--add finds homologous nucleotides in the query sequences that do not correspond to homologs in the backbone alignment, while UPP-add fails to recover these. This is an inherent limitation of HMM-based methods and motivates the development of EMMA.

## 2 New Method: EMMA

### 2.1 Comparing MAFFT--add and MAFFT-linsi--add

In a preliminary study (Experiment 0), we compared MAFFT--add and MAFFT-linsi--add to determine their relative accuracy and computational performance, especially as the number of sequences increased. We used 5000M2, a simulated dataset developed for this study (see Section 3.3.1), for this comparison. For this experiment, the backbone alignment has 1000 sequences and we varied the total number of added sequences from 100 to 2000.

**Figure 2:**
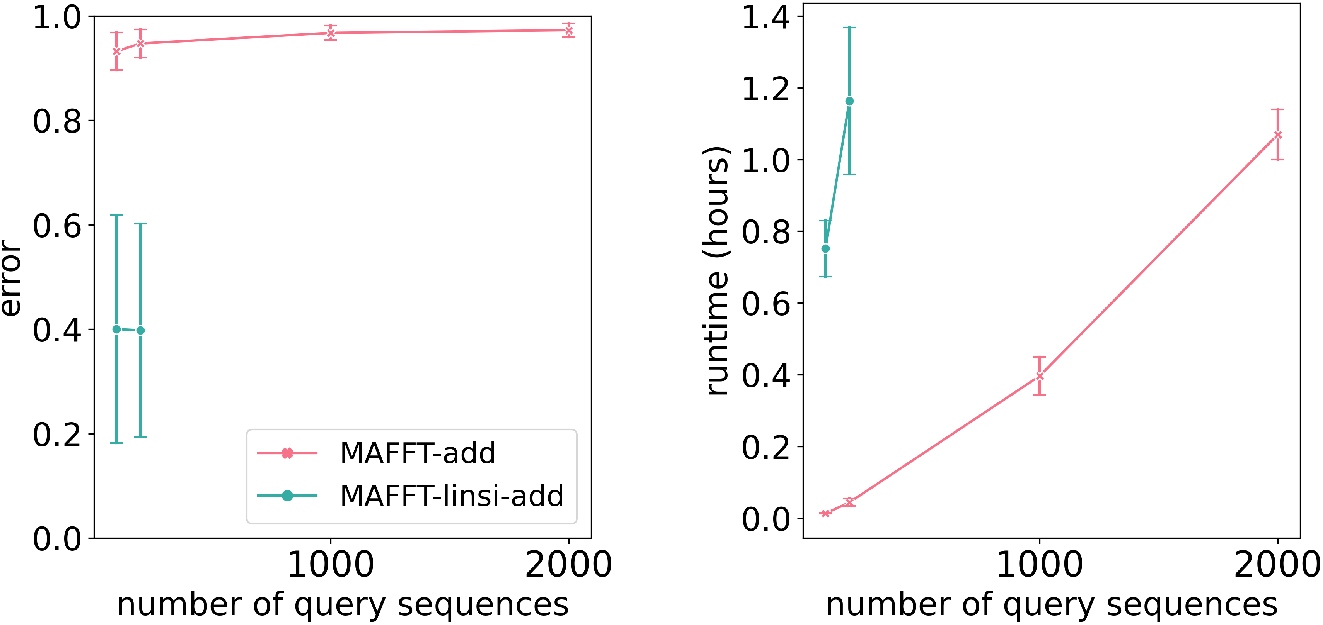
Experiment 0: MAFFT-linsi--add scalability issues. Alignment error (left) and runtime in hours (right) of MAFFT--add and MAFFT-linsi--add for adding 100, 200, 1000, or 2000 sequences to a 1000-taxon backbone alignment computed using MAGUS [13] on the INDELible 5000M2-het dataset with 5000 sequences. Averages over ten replicates are shown. Error bars shown for alignment errors are standard errors and standard deviations for runtime. We exclude replicate 4 because MAFFT-linsi--add encountered out-of-memory issues (64 GB) when adding 100 or 200 query sequences. Additionally, MAFFT-linsi--add is not shown for 1000 or 2000 query sequences because it either encountered out-of-memory issues or failed to complete within 12 hours. Alignment error is the average of SPFN (fraction of true pairwise homologies missing in the estimated alignment) and SPFP (fraction of pairwise homologies in the estimated alignment not found in the true alignment). Results for SPFN and SPFP separately show the same trends, and can be found in Supplementary Figure S1.

Figure 2 demonstrates that MAFFT-linsi--add (i.e., MAFFT--add run using -linsi) has a substantial accuracy improvement over MAFFT--add when run in default mode, but is limited to relatively small datasets. Hence, we are motivated to see if we can improve the scalability of MAFFT-linsi--add to large datasets.

### 2.2 EMMA’s algorithmic design

EMMA takes as input a multiple sequence alignment *C* on subset *S*_0_ and a set *S*_1_ of additional “query” sequences, and returns an alignment on *S* = *S*_0_ ∪*S*_1_ that is required to induce alignment *C* (thus *C* is treated as a constraint). We refer to *C* as a backbone alignment or as a constraint alignment.

EMMA was designed to enable MAFFT-linsi--add to scale to datasets where the total number of sequences in *S* (backbone plus query sequences) is very large. To achieve this, we use a divide-and-conquer strategy where we create a number of problems with smaller numbers of backbone sequences and query sequences on which MAFFT-linsi-add can efficiently and accurately run. We then show how we can combine the results into a solution to the overall problem of adding sequences into the backbone alignment. Some of the techniques in EMMA are adapted from PASTA [14] and UPP [4], as described below and in the supplementary materials.

EMMA has two algorithmic parameters, *l* and *u*, that govern the decomposition strategy; our study set default values for these parameters, but they can be supplied by the user. Given the input backbone alignment *C*, unaligned sequences, and (optional) values for parameters *l* and *u*, EMMA operates as follows:

**1. Step 1: Construct set of constraint subalignments**. Compute a tree on *C* (default: using the maximum likelihood heuristic FastTree2 [15]), and break it into smaller subsets by repeatedly deleting centroid edges (i.e., edges whose removal splits the leaf set into two sets of roughly equal size), and retaining only those subsets that contain at least *l* and at most *u* sequences. Each subset of the tree defines a subalignment *C*_*i*_ of *C* (defined by the set *S*_*i*_ of sequences in the subset). For each subalignment *C*_*i*_, *i* = 1 … *k*, construct a profile HMM.

**2. Step 2: Define the set of subproblems**. For each query sequence *q*, assign *q* to the constraint subalignment whose HMM has the best fit, as determined by the adjusted bitscore (introduced in [6]), which provides an estimate of the probability that the given HMM generates the given query sequence. This defines a set of subproblems: the constraint subalignment *C*_*i*_ and the set of query sequences *Q*_*i*_ assigned to *C*_*i*_. For each *i*, if the total number of sequences in *S*_*i*_ *∪ Q*_*i*_ exceeds 500, then partition the query sequences in *Q*_*i*_ into the smallest number of subsets needed in order to have at most 500 sequences in each subset, and ensure that have the subsets are as balanced in size as possible.

**3. Step 3: Apply MAFFT-linsi--add to add the query sequences**. For each resultant subproblem, use MAFFT-linsi--add to add all assigned query sequences to the selected constraint subalignment. The output of this step is a collection of extended subalignments (i.e., an alignment that agrees with the constraint alignment *C*_*i*_ but has some query sequences added).

**4. Step 4: Merge the extended subalignments using transitivity**. Each subalignment contains sites that come from the backbone alignment as well as newly introduced sites (representing homologies inferred between query sequence letters through the use of MAFFT-linsi--add). The sites from the backbone alignment allow us to merge these subalignments in an obvious way: if subalignments *A*_1_ and *A*_2_ both have sites drawn from site *j* from the backbone alignment, then these two sites are merged in the output alignment. Furthermore, the left-to-right ordering of columns in the subalignments can be used to define the left-to-right ordering in the output alignment. Transitivity was first used in PASTA and has been used in many subsequent methods, such as UPP, WITCH, and WITCH-ng. See Supplementary Materials Figure S2 for an example of transitivity merging.

#### Theorem 1.

*Given backbone alignment C on set S*_0_ *and query sequences S*_1_, *EMMA outputs an alignment on S*_0_ *∪ S*_1_ *that induces C when restricted to S*_0_.

*Proof*. Every subproblem consists of a constraint subalignment *C*^*′*^ and a set of query sequences *Q*^*′*^. MAFFT-linsi--add applied to subproblem (*C*^*′*^, *Q*^*′*^) is guaranteed to return an extended subalignment that induces *C*^*′*^ when restricted to the sequences for that set. Since every query sequence is in exactly one subalignment, all the extended subalignments will be consistent with *C*, the constraint alignment given to EMMA. By definition, the transitivity merge cannot undo homologies in any extended subalignment nor in the *i*. Furthermore, because each query sequence and each backbone sequence are in exactly one subalignment, the transitivity merge never merges columns in the constraint alignment; hence, it is guaranteed to return an alignment that induces the backbone alignment when restricted to the backbone sequences.

In addition to guaranteeing that the output of EMMA is consistent with the constraint alignment, the four-step design achieves several properties that are beneficial for alignment accuracy and scalability to a large number of sequences: (1) all runs of MAFFT-linsi--add have at most 500 sequences in total and (2) the division into subalignments is based on an estimated phylogeny, so that the sequences in each subalignment are likely to be closely related, and (3) each query sequence is assigned to a subalignment of sequences for which they are likely to be closely related (based on the bit-score calculation of the fit between the query sequence and the subalignment). These properties together make for subproblems that are small enough for MAFFTlinsi--add to run well on (i.e., we reduce the computational effort) and closely related enough for MAFFT-linsi--add to be highly accurate.

Note that EMMA uses HMMs to determine *which* subset alignment to assign a given query sequence but does not use the HMMs to perform the alignment. Instead, the alignment of the query sequence to the subalignment is performed using MAFFT-linsi--add in batch mode (i.e., all the query sequences assigned to the same subalignment are aligned together), which allows MAFFT-linsi--add to find homologies between letters in query sequences even if the backbone alignment does not have homologies as well.

## 3 Experimental design

### 3.1 Overview

Experiment 1 sets the parameters *l* and *u* in EMMA; these experiments are performed on the training data. In Experiment 2, we compare EMMA to MAFFT--add (v7.490, run in default mode), MAFFT-linsi--add (v7.490), and WITCH-ng-add (v0.0.2), using the testing datasets (disjoint from the training data).

All methods are evaluated for running time as well as alignment error (see below). Experiment 1 analyses were limited to 64 GB and 12 hours. Experiment 2 was run with 16 cores and 64 GB memory, with 24 hours of runtime. For MAFFT-linsi--add and MAFFT--add, we allowed 128 GB memory. All experiments were run on the UIUC Campus Cluster. See Supplementary Materials Section S3 for exact commands of all methods, and Supplementary Materials Section S4 for additional details on dataset generation.

### 3.2 Alignment error

For alignment error, we compare the estimated to the reference/true alignment, restricted to the (added) query sequences from the reference alignment. We report SPFN, SPFP, and expansion scores, defined as follows. The expansion score is the length of the estimated alignment divided by the length of the true or reference alignment; thus, the best result is 1.0, alignments that have expansion scores less than 1.0 are said to be “over-aligned”, while alignments that have expansion scores greater than 1.0 are “under-aligned”. The SPFN and SPFP error metrics are based on homologies, i.e., pairs of letters (nucleotides or amino acids) found in the same column in the true or estimated alignment. The sum-of-pairs false-negative (SPFN) rate is the fraction of homologies in the reference alignment that are missing in the estimated alignment. The sum-of-pairs false-positive (SPFP) rate is the fraction of pairs of homologies in the estimated alignment that are missing in the reference alignment. These metrics are calculated using FastSP [16].

These error metrics can be used to define accuracy measures in the usual way, with Recall the same as 1-SPFN and precision the same as 1-SPFP.

### 3.3 Datasets

We use both simulated and biological datasets of nucleotide and protein to evaluate EMMA. In addition to datasets from prior studies [17, 14, 18], we also generated one new simulated dataset using INDELible [19] and we use two new biological datasets (Rec and Res). The training datasets were used in Experiments 0 and 1, and all subsequent experiments used the testing data (a separate collection of datasets). Empirical statistics for these datasets can be found in Supplementary Materials Table S1. All datasets are described below, with the new datasets freely available online through the Illinois Data Bank (see Data Availability statement).

#### 3.3.1 Simulated datasets

All simulated datasets are based on evolving sequences down model trees with indels so that all sequence alignments involve gaps. As described below, these simulated datasets vary in terms of indel length distribution, rate of evolution, and whether the sites evolve identically and independently or under selection. All simulated datasets have at least 1000 sequences.

**ROSE** We included datasets from [17], which were generated using the ROSE [20] software. We used four model conditions (1000M1, 1000M2, 1000M3, and 1000M4) from [17]. Each model condition has 10 replicates, and each replicate has 1000 sequences, with an average sequence length of ∼1000 bp. The 1000M1–1000M4 models vary in rate of evolution (with 1000M1 the highest, and reducing rates as the index increases), thus enabling us to evaluate the impact of rates of evolution on alignment difficulty. All site evolution is non-ultrametric, and the sites evolve identically and independently.

**The INDELible simulated datasets** These datasets were generated for this study. We used INDELible [19] to evolve sequence datasets down a tree with a heterogeneous indel (insertion and deletion) model. Under this model, with a small probability, an indel event can be promoted to long indel events, modeling infrequent large gain or loss during the evolutionary process (e.g., domain-level indels). Hence, we name these new model conditions “het” to reflect their heterogeneous indels. Each replicate has 5000 sequences, and the model conditions range in evolutionary rates (and hence alignment difficulty), with model condition 5000M2-het having the highest rate of evolution, 5000M3-het somewhat slower, and 5000M4-het the slowest. We used non-ultrametric model trees, and all sites evolve identically and independently. See Supplementary Materials Section S4 for details of the data generation and Supplementary Materials Tables S2 and S3 for the parameter values used in the simulations.

**RNASim** The RNASim million-sequence dataset is from [14] and has been used in prior studies to evaluate alignment methods [4, 21]. In the RNASim simulation, RNA sequences evolve under a biophysical model and under selection in order to conserve the rRNA structure. Thus, the sites do not evolve independently. In this study, we subsampled 10 replicates, each with 10,000 sequences.

#### 3.3.2 Biological datasets

**CRW** The Comparative Ribosomal Website [18] (CRW) is a collection of nucleotide datasets with curated alignments based on secondary structure. We include 16S.3, 16S.T, and 16S.B.ALL, three large datasets from the CRW, with 5323, 6350, and 27643 sequences, respectively. These datasets have been used in previous studies [22, 4, 21] and exhibit sequence length heterogeneity. We use the cleaned versions from [21], for which any ambiguity codes or entirely gapped columns are removed. **10AA** The “10 AA” dataset is a publicly available collection of large curated protein alignments, originally assembled for the study evaluating PASTA [14], but also used to evaluate multiple sequence alignment methods [4, 22]. These curated alignments are based on protein structure and range from 320 to 807 sequences and include the eight largest BAliBASE datasets [23] and two datasets from [24].

**Serine Recombinases (Rec and Res)** These datasets were assembled for this study and are for two domains from serine recombinase. Protein sequences were taken from 350,378 GenBank bacterial and archaeal genome assemblies (Supplementary Materials Section S4) using Prodigal [25]. Serine recombinases were identified using the Pfam HMMs [26] Resolvase (Res) for the catalytic domain, and Recombinase (Rec) for the integrase-specific domain using *hmmsearch* from the HMMER package with the “trusted cutoffs” supplied by Pfam. Standards phiC31 and Bxb1 were added. From the 199,090 unique protein sequences, the Rec and Res domains were separately extracted using the boundaries determined by the HMM hits.

The Rec and Res datasets have reference alignments on a subset of the sequences (i.e., the seed alignment from Pfam), with Rec having 66 and Res having 112 seed sequences. Hence, the reference set for each dataset is much smaller than the entire set of sequences. Alignment error is evaluated only on these specific reference sequences.

#### 3.3.3 Constraint alignment selection

Recall that we have true alignments for the simulated datasets and reference alignments (based on structure) for the biological datasets. However, for the Rec and Res datasets, we have reference alignments only on a subset of the sequences. Alignment error is evaluated only on sequences for which we have reference alignments, which means that for the Rec and Res datasets we can only evaluate alignment error on a subset of the sequences.

In Experiment 2, we designed different ways of selecting a subset of the reference sequences to form the constraint alignment. The reference alignment, restricted to the selected sequences, is treated as the backbone alignment (i.e., the constraint alignment), and the remaining sequences are query sequences.

We considered three different scenarios for the selection of the backbone sequences, for which a constraint alignment would be provided by the user. The first two scenarios select sequences randomly from across the assume that the backbone sequences are selected randomly from the input. For these two scenarios, we vary the number of sampled sequences from “large” (at 25% of the full set of sequences) to “small” (at 10% of the sequences). The third scenario is designed for the case where a curated alignment is available for a small, and likely closely related set of sequences, found within a clade. .

##### 1. Large random subset

We begin by determining the set of reference sequences that are “full-length,” which means their length is within 25% of the median length. From the set of *F* full-length sequences, we randomly select min(1000, 0.25*F*) sequences.

##### 2. Small random subset

The protocol is identical to the large random subset protocol, except that we randomly select min(100, 0.1*F*) full-length sequences. However, if this number is less than 10, we just pick 10 sequences at random.

##### 3. Large clade-based subset

Here we restrict to sequences from a clade (as defined on either the true tree for simulated data or a maximum likelihood tree for the biological data), selecting a clade that has at most 1000 sequences, and that comes as close as possible to 25% of the reference sequences. In the case of ties, we pick the clade randomly,

## 4 Results

### 4.1 Experiment 1: Designing EMMA

Experiment 1 is the experiment where we set the algorithmic parameters, *l* and *u*, for EMMA. We vary *l* between 10 and 50 and *u* between *l* and 100, using the INDELiBLE 5000M2-het model condition, which has a high rate of evolution and so makes for difficult alignments. We use the *large random subset* scenario, so that the backbone alignment contains 1000 randomly selected sequences and the remaining 4000 sequences are query sequences.

Results for EMMA with different settings of (*l, u*) on 5000M2-het (10 replicates) are presented in Supplementary Materials Figure S3. The lowest error is found when *l* = 10 and that for this setting of *l, u* then has little impact. However, the setting for *u* does impact runtime, with the fastest runtime found (across all settings) with *l* = 10 and *u* = 25. Based on this experiment, we set these as the default settings for *l* and *u*.

### 4.2 Experiment 2: Evaluating EMMA to other sequence-adding methods

Here we show comparisons of EMMA using the default settings of *l* = 10 and *u* = 25 to WITCH-ng-add and MAFFT-linsi--add. We show a comparison to MAFFT—add run in default mode only for a limited set of analyses, since its error rates were much higher than the other methods; see Supplementary Materials Figures S4-S14 for full results including MAFFT--add.

**Figure 3:**
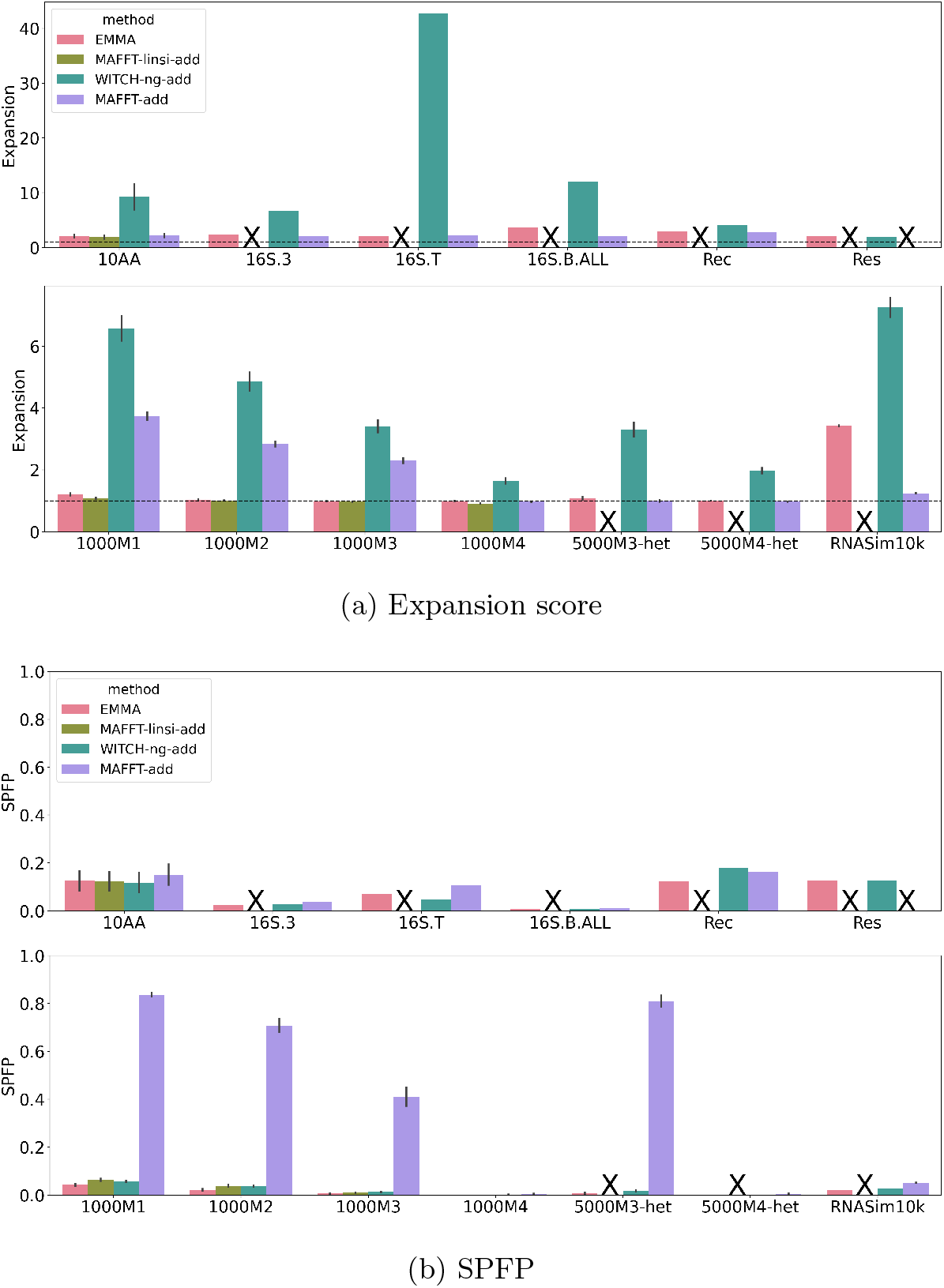
Expansion score (top) and SPFP (bottom) on large random backbones. In each subfigure, the top panel denotes biological datasets, and the bottom panel denotes simulated datasets (note the change in the y-axis range for expansion scores between the top and bottom panels). The horizontal dashed line indicates a perfect expansion score of 1. MAFFT-linsi--add failed to finish within 24 hours for datasets except for 10AA and ROSE 1000M1-4 and was not run on Rec and Res (due to their very large size, we knew it would not complete within the allowed time), and MAFFT--add encountered out-of-memory issues on the Res dataset (failed runs, or runs that were not attempted, are marked with “X”). The expansion score is the length of the estimated alignment normalized by the length of the reference or true alignment; optimal is 1.0, and values above 1.0 indicate alignments that are too long (and so are under-aligned).

#### 4.2.1 Expansion scores and SPFP

The expansion score is the estimated alignment length normalized by the reference or true alignment length. As seen in the Supplementary Materials Table S4, MAFFTlinsi--add had the best expansion scores, coming close to 1.0 in most cases. All the other methods, however, produced alignments that were longer than the true alignment (and hence were under-aligning), as indicated by expansion scores being greater than 1. MAFFT--add under-aligned the least of these methods, followed closely by EMMA, and WITCH-ng-add has by far the largest expansion scores. For the expansion score, WITCH-ng is the outlier, as its expansion scores were excessively high. This level of under-alignment also means that interpreting its SPFP rates is difficult since underalignment also reduces false positives.

With these observations, we now consider the SPFP scores. Results for large random backbones are provided in Figure 3, and results for the two other conditions are shown in Supplementary Materials Figures S4 and S5. WITCH-ng-add typically had among the best SPFP scores, but MAFFT-linsi-add and EMMA sometimes had better SPFP scores. Thus, the improved SPFP scores achieved by WITCH-ng-add are partially the result of its very high degree of under-alignment.

EMMA and MAFFT-linsi--add both had SPFP scores that are relatively close. There is an advantage to MAFFT-linsi--add over EMMA on the simulated ROSE datasets (1000M1–1000M4) but not on the 10AA datasets (the only biological datasets on which MAFFT-linsi--add completed).

We also see that MAFFT--add had higher SPFP scores than MAFFT-linsi--add on the datasets on which MAFFT-linsi--add could run. Given that MAFFT--add also had worse expansion scores than MAFFT-linsi--add, this indicates that MAFFT--add was generally inferior to MAFFT-linsi--add (consistent with Experiment 0, and also with other prior studies). Interestingly, although MAFFT--add had lower expansion scores than EMMA, it had higher SPFP scores. Given these trends, we omit MAFFT-add from the rest of the study (though see Supplementary Materials Figures S3–S8 for comparisons involving MAFFT--add).

We note that the SPFP scores, other than for WITCH-ng-add, were all relatively close (and generally low), making SPFP scores not very informative. Therefore, for the remainder of this study, we focus on SPFN, noting that 1-SPFN is the same as *recall* (or SP-Score).

#### 4.2.2 SPFN alignment error on large random backbones

Figure 4 shows the SPFN of the three sequence-adding methods (i.e., EMMA, WITCHng-add, and MAFFT-linsi--add) when adding to large random backbone alignments. The most striking observation is that MAFFT-linsi--add only succeeded in completing the datasets with at most 1000 sequences (i.e., the 10AA biological datasets and the ROSE simulated datasets with 1000 sequences); on all the other datasets, it failed to complete within the allowed time (24 hours). However, for those datasets that it completed, it had very good SPFN scores: better than the other methods for high rates of evolution (1000M1, 1000M2), in third place but still close to the best on the 10AA datasets, and tied for best on the ROSE datasets with low rates of evolution (1000M3 and 1000M4). We also see that EMMA was at least as accurate as WITCH-ng-add on nearly all datasets, except that WITCH-ng-add was better than EMMA on 1000M1.

**Figure 4:**
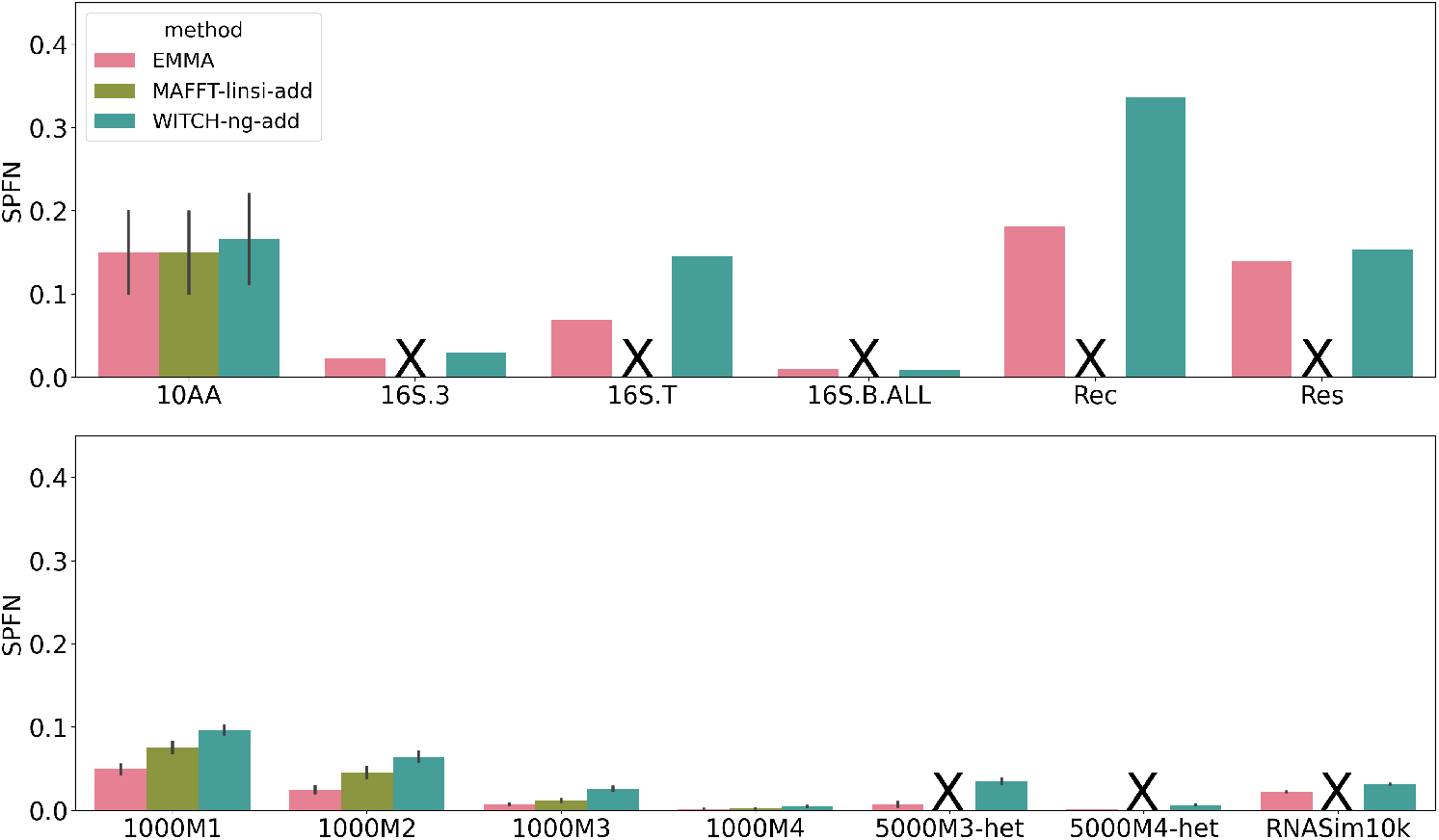
SPFN of EMMA, WITCH-ng-add, and MAFFT-linsi--add when adding sequences into large random backbone alignments. The top panel denotes biological datasets, and the bottom panel denotes simulated datasets. Error bars indicate standard errors. MAFFT-linsi--add only completed within the allowed 24 hours on datasets with at most 1000 sequences, and we did not run it on Rec or Res due to their large numbers of sequences (failed runs, or runs that were not attempted, are marked with “X”). Comparisons with MAFFT--add can be found in Supplementary Figure S7. SPFN results for each of the 10AA datasets are in Supplementary Figure S12.

#### 4.2.3 SPFN alignment error on small random backbones

As with large random backbones, MAFFT-linsi--add only succeeded in finishing on the datasets with at most 1000 sequences (Figure 5), but on those datasets, it was among the most accurate methods. The relative accuracy between the three methods is very similar to that displayed on the large backbones so EMMA was generally at least as accurate as WITCH-ng-add and, in some cases, was much more accurate, while MAFFT-linsi--add had the best accuracy of all methods when it can run). Finally, all methods had somewhat larger error rates in this condition than when placed into a large random backbone.

**Figure 5:**
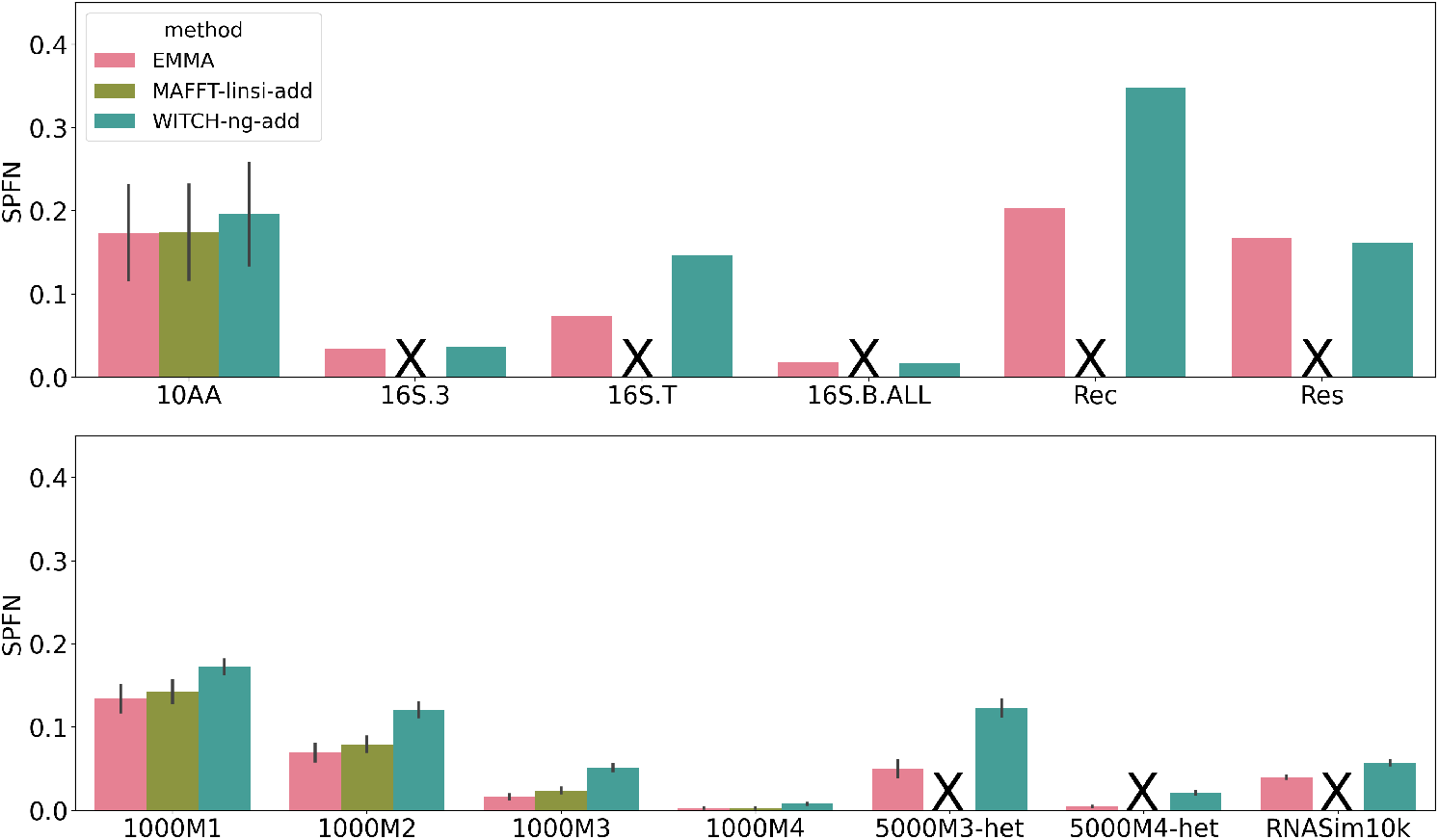
SPFN of EMMA, WITCH-ng-add, and MAFFT-linsi--add when adding sequences into small random backbone alignments. The top panel denotes biological datasets, and the bottom panel denotes simulated datasets. Error bars indicate standard errors. MAFFT-linsi--add failed to finish within 24 hours for datasets except for 10AA and ROSE 1000M1-4, and we did not run it on Rec or Res due to their large numbers of sequences (failed runs, or runs that were not attempted, are marked with “X”). Comparisons with MAFFT--add can be found in Supplementary Figure S8. SPFN results for each of the 10AA datasets are in Supplementary Figure S13.

#### 4.2.4 SPFN alignment error on clade-based backbones

As with the other conditions, when adding sequences into a clade-based backbone alignment, MAFFT-linsi--add failed to complete on any dataset with more than 1000 sequences. However, in many respects, the results for the clade-based backbone are different than for the random backbones (Figure 6). The most noteworthy difference is that error rates increased for all methods when given clade-based backbones instead of random backbones, but the increase was largest for WITCH-ng-add. The gaps between methods increased, with WITCH-ng-add no longer tying for best in any condition. MAFFT-linsi--add is clearly the most accurate of all the methods on the 1000M1, 1000M2, and 1000M3 datasets, and then ties for best on 1000M4. EMMA is strictly more accurate than WITCH-ng-add, usually by a large margin, on all datasets. Nevertheless, both EMMA and WITCH-ng-add have high errors on datasets with high rates of evolution (1000M1, 5000M3-het).

**Figure 6:**
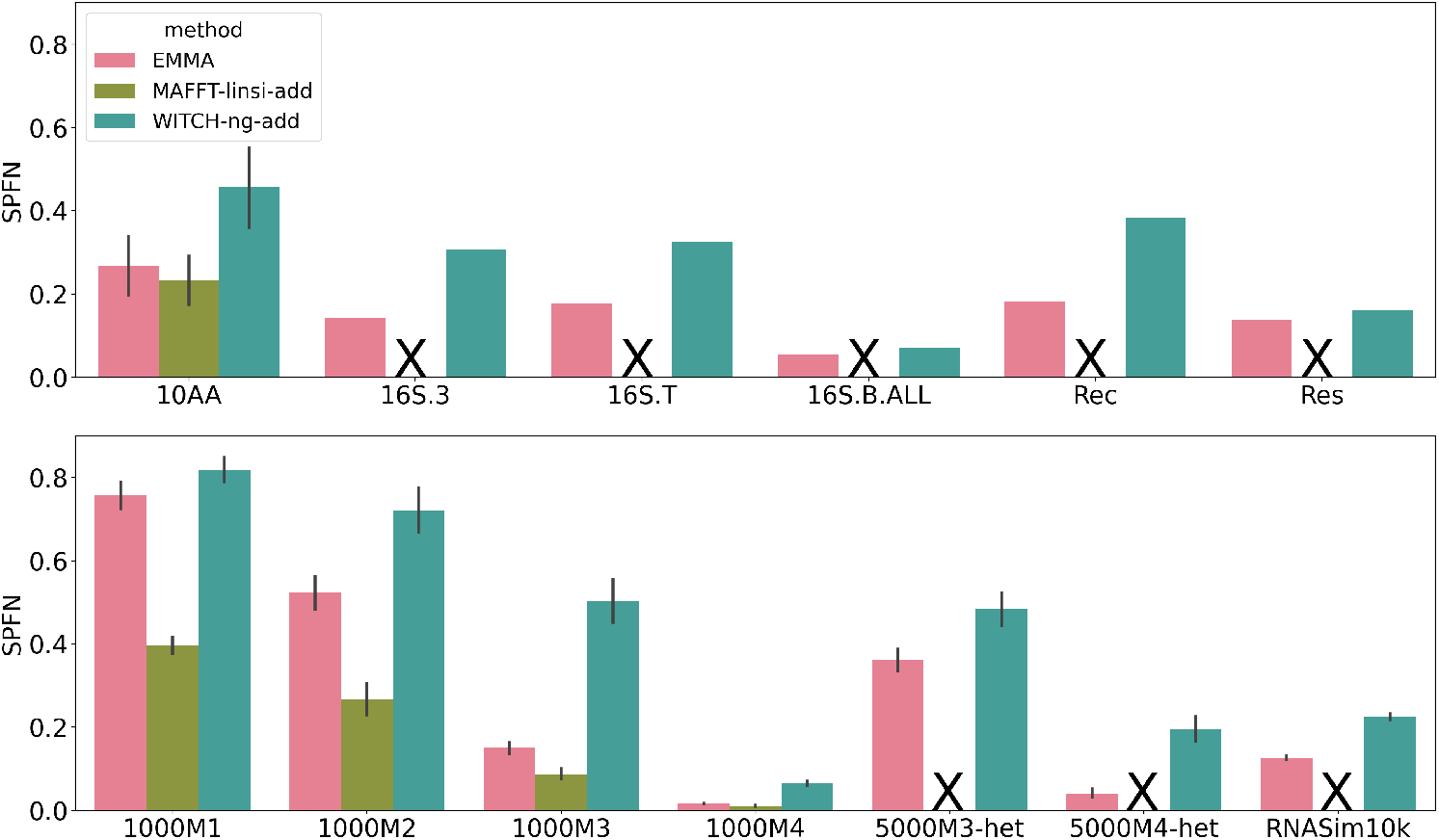
SPFN of EMMA, WITCH-ng-add, and MAFFT-linsi--add when adding sequences into clade-based backbone alignments. The top panel denotes biological datasets, and the bottom panel denotes simulated datasets. Error bars indicate standard errors. MAFFT-linsi--add failed to finish within 24 hours for datasets except for 10AA and ROSE 1000M1-4, and we did not run it on Rec or Res due to their large numbers of sequences (failed runs, or runs that were not attempted, are marked with “X”). Comparisons with MAFFT--add can be found in Supplementary Figure S9. SPFN results for each of the 10AA datasets are in Supplementary Figure S14.

#### 4.2.5 Computational performance

MAFFT-linsi--add and MAFFT--add are the only methods that failed to complete on at least one of the datasets within the allowed runtime (24 hours), using 16 cores and 128 GB memory. Supplementary Materials Section S5 shows that both methods reported out-of-memory issues or crashes. Specifically, MAFFT-linsi--add had an outof-memory issue when attempting to analyze the INDELible 5000M2-het training dataset and allowed 64 GB. We also noted that MAFFT--add had an out-of-memory issue on the Res dataset with ∼186K sequences, even when allowed 128 GB of memory. This is perhaps not surprising, since MAFFT--add also computes the *n ×n* pairwise distance matrix, where *n* is the number of sequences in total (backbone sequences and query sequences together), and shows that MAFFT--add as well as MAFFT-linsi--add both have large memory requirements.

Figure 7 shows a comparison between the methods given a large random backbone. MAFFT-linsi--add was the slowest method on all the datasets when it could run (as noted above, it failed to complete on any dataset with more than 1000 sequences). WITCH-ng-add was overall the next slowest method, but it was faster than EMMA on the 10AA datasets (which are the smallest datasets we examined, all below 1000 sequences). MAFFT--add was overall the fastest method, but WITCH-ng-add was faster on the Rec dataset and EMMA was faster on the 5000M3-het dataset. Overall, EMMA was in between WITCH-ng-add and MAFFT--add in terms of running time on these datasets.

**Figure 7:**
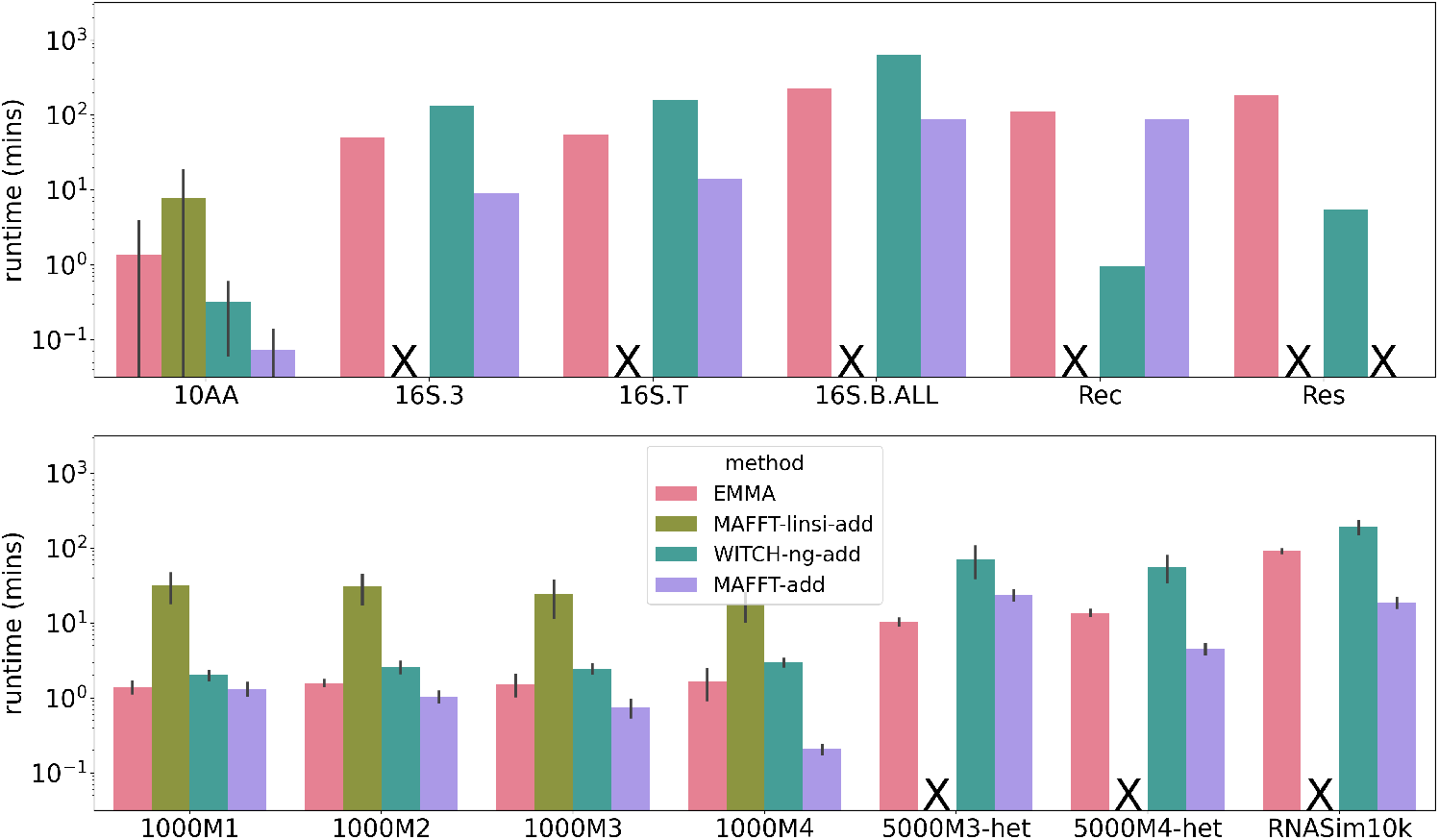
Experiment 2: Runtime (log-scale) in minutes when adding to large random backbone alignments. The top panel denotes biological datasets, and the bottom panel denotes simulated datasets. Except for the datasets with at most 1000 sequences, MAFFT-linsi--add failed to finish within 24 hours and was not attempted on Rec and Res due to their large number of sequences. MAFFT--add encountered out-of-memory issues on the Res dataset. Failed runs, or runs not attempted due to dataset size, are marked with “X”. Error bars indicate standard deviation.

The runtimes of all methods on the other conditions (small random backbones and clade-based backbones) are shown in Supplementary Materials Figures S10 and S11. On these datasets, we see somewhat similar trends, with EMMA faster than MAFFTlinsi--add on the datasets where MAFFT-linsi--add can run, but now EMMA is slower than both WITCH-ng-add and MAFFT--add.

The only methods that succeeded in completing all datasets were EMMA and WITCH-ng-add, and the maximum runtimes used by EMMA and WITCH-ng-add were 12.2 and 10.5 hours, respectively. Given that WITCH-ng-add and EMMA completed so quickly using under 64 GB memory on these datasets and that the datasets had up to ∼186,000 sequences, it is reasonable to say that WITCH-ng-add and EMMA are acceptable with respect to speed and scalability.

We also tried to give MAFFT-linsi--add three days (72 hours instead of 24 hours) to run on 16S.3 and 16S.T for the small random backbone and large clade-based backbone, but it still was not able to complete. Finally, we consider MAFFT--add to be reasonably scalable since it only had computational problems with the largest dataset (Res), and it is by far the fastest method we tested.

## 5 Discussion

The four methods we have evaluated – EMMA, MAFFT--add, MAFFT-linsi--add, and WITCH-ng-add, vary in their ability to run on large datasets (i.e., scalability), runtime, and accuracy (here focused on SPFN). The relative performance is summarized in Table 1, and discussed further below.

**Table 1:** Overall performance of the tested methods. For scalability, we indicate which study datasets the method completed on. For SPFN and speed, we indicate the relative position (1 is best, 2 is second best, etc.).

| Criterion | EMMA | WITCH-ng-add | MAFFT--add | MAFFT-linsi--add |
| --- | --- | --- | --- | --- |
| Scalability | all | all | all but largest | $\leq 1000$ sequences |
| SPFN-random | 1 | 2 | 3 | 1 |
| SPFN-clade | 2 | 3 | 4 | 1 |
| Speed | 2 | 2 | 1 | 3 |

Our study showed clear differences in scalability, with EMMA and WITCH-ng-add the most scalable (completing on all datasets within the time and memory limits), MAFFT--add the next most scalable (failing to complete on one dataset), and MAFFT-linsi--add the least scalable (failing on all but the datasets with at most 1000 sequences). It is easy to see why EMMA can scale to the largest datasets, as the decomposition strategy and transitivity merge are both very fast and the only potentially computationally intensive part is when it runs MAFFT-linsi--add on subsets of query sequences to add them to subsets of the backbone alignment. However, by design, the subproblems are all small (at most *u* = 25 backbone sequences and at most 500 query sequences). This design strategy allows EMMA to complete on all the datasets in our study. It is also easy to see why MAFFT-linsi--add is limited in scalability since its algorithmic design involves a quadratic runtime.

Accuracy differences were also apparent, but were generally very small when the rates of evolution were low enough. This also is as expected, since prior studies have shown differences in alignment error often are very small or negligible when sequence identity is high, which occurs under low rates of evolution (e.g., [27, 28]).

When comparing accuracy under higher rates of evolution, we see that the selection strategy for the backbone sequences – i.e., whether they were selected randomly or from within a clade, and the number of backbone sequences – impacts both absolute and relative alignment error. In particular, all methods have better accuracy with random sampling than with clade-based sampling, and also have better accuracy when there are more sequences in the backbone alignment. These trends are expected, since the constraint alignment is used as a model of the family and dense sampling throughout the family (i.e., the large random backbone) ensures a better model than sparse random sampling (i.e.,, the small random backbone) or sampling from just within a clade.

The comparison between EMMA and WITCH-ng-add with respect to accuracy favors EMMA across all sampling strategies, although the two methods have nearly identical accuracy for some conditions. The accuracy advantage of EMMA is more noticeable on datasets with high rates of evolution, such as ROSE 1000M1, 1000M2, and INDELible 5000M3-het. Since WITCH-ng-add relies on using HMMs created from the backbone to align the queries whereas EMMA uses MAFFT-linsi--add to align the queries, the improvement in accuracy for EMMA is likely to be due to the greater sensitivity of MAFFT-linsi--add for detecting homologies than the HMM-based approach within WITCH-ng-add.

The comparison between MAFFT-linsi--add and EMMA, although restricted to just those datasets on which MAFFT-linsi--add could run (i.e., the datasets with at most 1000 sequences), reveals the following trends. When given a large random backbone, EMMA matches or improves on MAFFT-linsi--add under all conditions, but has an advantage given on the ROSE 1000M1 and 1000M2 conditions, and then matches MAFFT-linsi--add for the 1000M3 and 1000M4 conditions. The same relative performance is seen for the small random backbone, but with a smaller difference between the methods. Thus, the rate of evolution impacts the relative accuracy of the two methods, favoring EMMA when the rate of evolution is high, suggesting that MAFFT-linsi--add accuracy is impacted more substantially by the rate of evolution than EMMA. This trend is consistent with prior studies showing that MAFFT-linsi is sensitive to rate of evolution, an observation that led to the design of SATé [27] and its descendent methods [29, 14, 13].

Interestingly, MAFFT-linsi--add improves on EMMA when the sequences are drawn from a clade, with a substantial improvement under the higher rate of evolution. In other words, when the backbone is drawn from a clade, the algorithmic strategy in EMMA – which applies MAFFT-linsi--add to subsets that contain at most 25 backbone sequences from the clade – is inferior to using the entire backbone. This is clearly a condition where the EMMA divide-and-conquer strategy is not well-suited.

In understanding the conditions where MAFFT-linsi--add is more accurate than EMMA, it is helpful to understand that the homologies between query sequences found by EMMA are either found directly by MAFFT-linsi-add on a small subset (with at most 500 sequences) or inferred when combining these extended alignments using transitivity. Thus, the relative accuracy of EMMA and MAFFT-linsi--add is impacted by the divide-and-conquer strategy, and in particular the size of the subsets in the decomposition of the backbone tree. Given that our default setting currently sets *u* = 25, this choice creates subsets that have very few backbone sequences (at most 25) and then adds query sequence, but not so many as to exceed 500 total sequences. This is potentially why MAFFT-linsi-add can be more accurate than EMMA, since MAFFT-linsi-add can find homologies between all pairs of query sequences, whereas EMMA (which applies MAFFT-linsi-add to small subsets) can only achieve this within the subsets it produces. Our Experiment 1, where we set the parameters *l* = 10 and *u* = 25, was based on the INDELible 5000M2 dataset, which has a high rate of evolution. It seems likely that other settings for *l* and *u* might lead to improved accuracy when the model condition has a lower rate of evolution, since MAFFT-linsi--add’s accuracy improves as the rate of evolution decreases. In other words, EMMA’s algorithmic design aimed to reduce runtime and improve scalability, but for some challenging inputs (e.g., clade-based backbones) where MAFFT-linsi--add can run, the algorithmic design of EMMA may reduce accuracy relative to MAFFT-linsi--add. Hence, there is room for improvement in designing EMMA.

However, MAFFT-linsi--add could not complete on the study datasets with more than 1000 sequences given the limitations in computational resources we imposed. In contrast, EMMA was able to complete on all study datasets. Thus, while MAFFTlinsi--add had a clear accuracy advantage over EMMA for the clade-based backbones (though not on the random backbones), EMMA has a computational advantage over MAFFT-lines--add in being able to scale to larger datasets.

Given the impact of clade-based sampling for the backbone sequences, it is important to remember that EMMA is never identical to MAFFT-lines--add, even when analyzing small subsets. When we run MAFFT-linsi--add on the datasets with at most 1000 sequences, all the query sequences are considered together when they are added into the backbone alignment. In contrast, by construction EMMA only adds query sequences to subsets of the backbone alignment with at most *u* sequences, and in our study we set *u* = 25 by default. From a computational standpoint, this strategy allows EMMA to scale to very large datasets. From an accuracy perspective, however, the impact clearly depends on how the backbone sequences were sampled. EMMA has an accuracy advantage over MAFFT-linsi--add when the backbone sequences are randomly sampled and there is a high rate of evolution, and matches MAFFT-linsi--add for accuracy when the backbone sequences are randomly sampled but there is a lower rate of evolution. However, as we have seen, this strategy is poor when the backbone sequences are sampled from a clade and the rate of evolution is high.

This observation also indicates that two-phase multiple sequence alignment methods that operate by selecting and aligning a set of sequences for the backbone, and then add in the remaining sequences, should – if possible – select the backbone sequences from across the evolutionary tree for the dataset, in order to obtain the best representative model of the gene family.

## 6 Conclusions

This study presented EMMA, a new method for adding sequences into existing constraint alignments (also called backbone alignments). EMMA uses a divideand-conquer strategy to enable MAFFT-linsi--add, a highly accurate version of MAFFT--add, to scale to large datasets. By itself, MAFFT-linsi--add was unable to complete using the computational resources on the datasets we studied with more than 1000 sequences, but EMMA succeeded in completing with even fewer resources on the largest dataset we studied, with more than 180,000 sequences.

Our study shows that EMMA has comparable or better alignment accuracy than MAFFT--add and WITCH-ng-add under all the conditions tested, and also comparable or better accuracy than MAFFT-linsi--add when the backbone sequences are selected randomly. However, our study also showed that when the backbone sequences are selected from a clade and the dataset is sufficiently small, then MAFFT-linsi--add is more accurate than EMMA. This finding indicates clearly that the design of EMMA needs to be reconsidered for the case where the user provides a curated alignment for a closely related group of sequences contained within a clade, and wishes to add additional sequences that are distantly related to the backbone sequences.

Thus, future work should investigate appropriate modifications to the divide-and-conquer strategy in EMMA to address the case where the backbone sequences are distantly related from the query sequences. EMMA alignment accuracy could potentially be improved with a re-alignment stage to see if sets of sites could be merged together through the detection of additional homologies between query sequences. We should also explore other biological datasets to document the accuracy of these methods under a wider range of real-world conditions. EMMA is not yet optimized for speed, and a more careful parallel implementation may provide a substantial speedup. Finally, EMMA could be studied as the second stage of the standard two-stage pipeline protocol used by UPP/UPP2, WITCH, and WITCH-ng for *de novo* multiple sequence alignment. Given the high accuracy of these two-stage methods for aligning datasets with high sequence length heterogeneity, this is likely to provide improved accuracy.

## Ethics approval and consent to participate

NA

## Consent for publication

NA

## Availability of data and materials

The INDELible, Rec, and Res datasets were created for this study. The Rec and Res datasets are available in the Illinois Data Bank at https://doi.org/10.13012/B2IDB-2567453_V1. The subsampled RNASim datasets (10,000 sequences for each replicate) are available at https://doi.org/10.13012/B2IDB-4194451_V1. Pfam seed sequences are included for both datasets (66 and 112 sequences for Rec and Res, respectively). The INDELible datasets (5000M2-het, 5000M3-het, and 5000M4-het) are available at https://databank.illinois.edu/datasets/IDB-3974819.

All other datasets are from prior studies and are available in public repositories. The 10AA datasets are available at https://sites.google.com/eng.ucsd.edu/datasets/alignment/pastaupp. The three CRW datasets are available at https://databank.illinois.edu/datasets/IDB-2419626. The ROSE datasets are available at https://sites.google.com/eng.ucsd.edu/datasets/alignment/sate-i.

## Author’s contributions

CS developed the EMMA software, performed the analyses, interpreted the data, and drafted the manuscript. BL provided simulated datasets. KPW provided biological datasets and interpreted the data.. TW conceived of the approach, designed the study, interpreted the data, wrote the final draft, and supervised the research. All authors read and approved the final manuscript.

