## Supplementary Materials for "EMMA: A New Method for Computing Multiple Sequence Alignments given a Constraint Subset Alignment"

### Supplementary Materials for "Computing Multiple Sequence Alignments given a Constraint Subset Alignment using EMMA"

**S5 Additional Figures** **14**

**S6 Additional Tables** **28**

S4 Experiment 2: Expansion and SPFP on large random backbones. 17
S5 Experiment 2: Expansion and SPFP on small random backbones. 18
S6 Experiment 2: Expansion and SPFP on clade-based backbone . . 19
S7 Experiment 2: SPFN on large random backbones . . . . . . . . . 20
S8 Experiment 2: SPFN on small random backbones. . . . . . . . . 21
S9 Experiment 2: SPFN on clade-based backbones. . . . . . . . . . 22
S10 Experiment 2: Runtime on small random backbones. . . . . . . . 23
S11 Experiment 2: Runtime on clade-based backbones. . . . . . . . . 24
S12 Experiment 2: Individual SPFN for 10AA datasets on large random backbones. . . . . . . . . . . . . . . . . . . . . . . . . . . . . 25
S13 Experiment 2: Individual SPFN for 10AA datasets on small random backbones. . . . . . . . . . . . . . . . . . . . . . . . . . . . . 25
S14 Experiment 2: Individual SPFN for 10AA datasets on clade-based backbones. . . . . . . . . . . . . . . . . . . . . . . . . . . . . 26

### S1 Additional information about prior methods

#### S1.1 UPP-add

UPP [Nguyen et al., 2015] is a method that was designed to align sequence datasets that have high sequence length heterogeneity. UPP uses two stages, where the first stage extracts and aligns a subset of the sequences deemed to be full-length, and the second stage adds in the remaining sequences into the alignment from the first stage (which we refer to as the "backbone alignment"). In this paper, we focus on the second stage of UPP, which we will refer to as UPP-add.

Given a backbone alignment, UPP-add builds an ensemble of Hidden Markov Models (i.e., an eHMM) to represent the backbone alignment, which it then uses to add query sequences into the backbone alignment. To build the eHMM, UPP-add computes a backbone tree on the backbone alignment using the maximum likelihood heuristic FastTree2 [Price et al., 2010]. Then, UPP-add recursively decomposes the backbone tree into smaller subtrees by deleting "centroid edges" until the last decomposition results in subtrees with at most $A$ leaves ($A = 10$ in default UPP). Each subtree thus defines a subset of the backbone sequences and hence also a subalignment (induced by the backbone alignment on the subset of sequences). For each subalignment, an HMM is created using *hmmbuild* from the HMMER suite [Finn et al., 2011]. Each query sequence is searched against all HMMs in the eHMM using *hmmsearch* (also from HMMER), and then mapped to the HMM with the highest bit-score. The alignment of the query sequence to the subalignment for the selected HMM is computed using *hmmalign* (also from HMMER). Because the subalignments are induced by the backbone alignment, this also defines an alignment of the query sequence to the backbone alignment, which we refer to as an "extended alignment". The set of all such extended alignments (one for each query sequence) are merged using transitivity to form the final alignment.

#### S1.2 WITCH-add and WITCH-ng-add

WITCH [Shen et al., 2022] and WITCH-ng [Liu and Warnow, 2023] are two MSA methods that use the same type of two-stage approach as UPP, where the first stage extracts and aligns a subset of the sequences, and the second stage adds the remaining sequences into the backbone alignment. Here we focus on the second stages of these methods, i.e., WITCH-add and WITCH-ng-add.

Like UPP-add, both WITCH-add and WITCH-ng-add compute the backbone tree and eHMM, and then perform an all-against-all search between all query sequences and all HMMs. Then, for a given query sequence $q$, they compute a weight between for each query-HMM pair based on an "adjusted bitscore", as described in Shen et al. [2022], and select the top $k$ HMMs for that query sequence ($k = 10$ in default WITCH). As described in UPP-add, each such query-HMM pair defines a specific extended alignment for the query sequence, i.e., a specific way of adding the query sequence to the backbone alignment. WITCH-add and WITCH-ng-add each compute a consensus of these $k$ extended alignments, and differ in how these consensus alignments are computed. WITCH-add computes a consensus of these $k$ extended alignments using the Graph Clustering Merger (GCM), a technique from MAGUS [Smirnov and

Warnow, 2021], while WITCH-ng-add uses a simple variant of Smith-Waterman [Smith and Waterman, 1981] to compute the consensus alignment. These consensus alignments thus define a single extended alignment for the query sequence to the backbone alignment. Finally, as in UPP-add, these extended alignments are merged together using transitivity. WITCH-add in general, has better alignment accuracy than UPP-add [Shen et al., 2022], but WITCH-ng-add is faster than WITCH and at least as accurate [Liu and Warnow, 2023]. Thus, in this study, we examined WITCH-ng-add but not WITCH-add.

#### S1.3 Transitivity Merger

Transitivity is a mathematical property for some binary relations. It says that if $X$ and $Y$ are related, and if $Y$ and $Z$ are related, then $X$ and $Z$ must also be related. Since homology (i.e., descent from a common ancestor) is such a relationship, and since the objective of multiple sequence alignment is to detect homology, we will be able to use transitivity to compatible alignments that share some letters through transitivity.

The Transitivity Merger is used in any two-phase method that constructs an alignment on a subset and then adds each of the remaining sequences to the subset alignment. The collection of extended alignments (one such alignment for each remaining sequence) must be merged together, and the merging process is produced using transitivity. Thus, it is a fundamental step in UPP Nguyen et al. [2015]. However, it is also used in PASTA Mirarab et al. [2015], which produces disjoint alignments, then merges pairs of these alignments, producing overlapping subsets that are compatible with each other. To merge this set of overlapping alignments, it uses transitivity. Thus, it is also a fundamental step in PASTA.

In Figure S2 we provide an example of how transitivity is used to merge two alignments, in the context of a two-phase method where the backbone alignment is divided into two subsets. The backbone alignment has four sequences and is split into two subalignments, each with two sequences. Then, two query sequences `AACTA` and `AATCAA` are added to these two subalignments, producing two extended subalignments. Since the two subalignments were taken from the backbone alignment, each site in each extended subalignment corresponds to either a specific site in the backbone alignment or corresponds to an insertion. In general, insertions can be between sites, but in the example shown, each insertion is at the end of the subalignment. We show how this information is used to merge the two extended subalignments.

The first `A` in query sequence `AACTA` is aligned to the first column of the first subalignment, which implies it will be placed in the first column of the final alignment, which will also contain the nucleotides from the first column of the backbone alignment. This is an application of transitivity.

The same argument shows that the first `A` in query sequence `AATCAA` is aligned to the first column of the second subalignment, which implies it *also* will be placed in the first column of the backbone alignment. Thus, the first `A`s in these query sequences will be considered homologous to each other, as well as to all the nucleotides in the first column of the backbone alignment, and placed in the first column of the final alignment. This is also an application of transitivity.

In contrast, the last `A` in each of the two query sequences is placed in the last column by itself within their extended subalignments. This means each of these

As are not considered homologous to any letter in their extended subalignments, and hence, they cannot be in a column with any other letter in the final alignment. Hence, the final alignment must have two final columns, each containing one of these two final As and no other letters (otherwise, some homology would be implied). However, the order of these two final columns is not determined, so the transitivity merge allows both outcomes.

#### S1.4 Adjusted Bitscore

The adjusted bitscore is given by the following equation:

$$w_{H_i,q} = P(H_i|q) = \frac{1}{\sum_{j=1}^{d} 2^{BS(H_j,q)-BS(H_i,q)+\log_2 \frac{s_j}{s_i}}} \quad (1)$$

where $H_i$ represents a hidden Markov model (HMM), $q$ is a query sequence to be added, $d$ is the total number of HMMs, $BS(H_i, q)$ is the bitscore between $H_i$ and $q$ (computed by HMMER [Finn et al., 2011]), and $s_i$ is the number of sequences in the alignment that generates $H_i$. For additional details, see Shen et al. [2022], Section 2.4.

#### S2 Commands for software

All commands are based on the assumption that 16 CPU cores are available.

##### S2.1 Backbone generation and sequence-adding methods

1. For Experiment 0 only, we computed a MAGUS backbone alignment (GitHub version committed on April 5th 2021):

```
$ python3 magus.py --recurse false -np 16  \
     -i [unaligned backbone sequences] -d [outdir] \
     -o [output alignment]
```

2. We used FastTree2 (v2.1 multi-threaded version) to generate a backbone tree from a given constraint (backbone) alignment for each dataset (`-nt -gtr` for nucleotide or `-lg` for amino acids):

```
$ FastTreeMP {-nt -gtr|-lg} [backbone alignment] > [backbone
    tree]
```

3. We used RAxML-ng (v1.0.3) to generate the reference tree for Rec and Res datasets (for selecting a clade of sequences in Experiment 2 - adding to a clade-based backbone). We ran bootstrapping on their reference alignments and used the first tree returned:

```
$ raxml-ng --bootstrap --msa [reference alignment] --model LG+
    G \
    --bs-trees 500 --force perf_threads \
    --threads 16 --prefix [output prefix]
```

4. WITCH-ng-add (v0.0.2) to add query sequences to an existing backbone alignment and tree:

```
$ witch-ng add -i [query sequences path] -b [backbone
    alignment] \
    -t [backbone tree] -o [output alignment] --threads 16
```

5. MAFFT/MAFFT-linsi --add (MAFFT v7.490) to add query sequences to an existing backbone alignment:

```
$ [mafft/mafft-linsi] --quiet --thread 16 --add [query
    sequences] \
    [backbone alignment] > [output alignment]
```

##### S2.2 Evaluation

1. FastSP (v1.7.1) to obtain SPFN and SPFP. We average these two results to produce an overall "Alignment error". The "-ml -mlr" options will ignore columns with lowercase letters (considered as insertions in UPP and WITCH):

```
$ java -jar FastSP.jar -ml -mlr -e [estimated alignment] \
    -r [reference alignment] -o [output file]
```

2. Runtime is obtained by adding the following command before any software:

```
$ { /usr/bin/time -v [software command and options] ; } \
    > [runtime information]
```

3. Empirical dataset statistics (i.e., p-distance, % gaps, etc.) are obtained using the following script (https://wiki.illinois.edu/wiki/display/warnowlab/Get+alignment+statistics). Alternatively link from Dropbox (https://www.dropbox.com/s/wqt8hpvqkdjge04/alignment_stats.zip?dl=0):

```
$ java -cp ./alignment_stat AlignmentStatistics \
    [alignment file] [output file]
```

### S3 Datasets

#### S3.1 Empirical statistics

Table S1 provides empirical statistics for the datasets used in this study. We include both nucleotide (NT) and protein (AA) datasets. Datasets marked with (*) denote training datasets. The number of replicates (if more than one) is specified with a parenthesis after the dataset name.

The p-distance is defined as follows. The Hamming distance between two (aligned) sequences is defined as the number of columns that have different letters and neither is gapped. The normalized Hamming distance is the Hamming distance divided by the total number of columns where both sequences have letters, thus producing values between 0.0 and 1.0. This value is known as the p-distance.

The percent gaps is the percentage of the alignment that is occupied by "-".

The Indelible, ROSE, and RNASim datasets are all simulated, and the remaining sequence datasets are biological. Except for Rec and Res, there are either true (known because simulated) or curated alignments on all the sequences in the dataset. The Rec and Res datasets have curated alignments only on a subset of the sequences (66 for Rec and 112 for Res). See the main paper for additional details.

**Table S1: Empirical statistics of study datasets. "*" marks the training dataset. Numbers in parentheses after dataset names indicate the number of replicates.**

| **Dataset** | **# Seqs** | **p-distance** | | **% gaps** | **Avg. Seq. length** | **Align length** |
|---|---|---|---|---|---|---|
| | | **Avg.** | **Max** | | | |
| Indelible datasets (NT) | | | | | | |
| - 5000M2-het(10)* | 5000 | 0.686 | 0.789 | 97.9 | 1025 | 48,887 |
| - 5000M3-het(10) | 5000 | 0.664 | 0.764 | 96.0 | 1001 | 25,262 |
| - 5000M4-het(10) | 5000 | 0.528 | 0.671 | 96.0 | 1001 | 24,415 |
| ROSE datasets (NT) | | | | | | |
| - 1000M1(10) | 1000 | 0.696 | 0.768 | 74.2 | 1011 | 3934 |
| - 1000M2(10) | 1000 | 0.681 | 0.761 | 73.6 | 1014 | 3896 |
| - 1000M3(10) | 1000 | 0.658 | 0.741 | 62.3 | 1007 | 2688 |
| - 1000M4(10) | 1000 | 0.496 | 0.607 | 60.6 | 1008 | 2575 |
| RNASim dataset (NT) | | | | | | |
| - RNASim10k(10) | 10,000 | 0.411 | 0.616 | 82.2 | 1551 | 8694 |
| *CRW* datasets (NT) | | | | | | |
| - 16S.3 | 6323 | 0.315 | 0.833 | 82.1 | 1557 | 8716 |
| - 16S.T | 7350 | 0.345 | 0.901 | 87.4 | 1492 | 11,856 |
| - 16S.B.ALL | 27,643 | 0.210 | 0.769 | 79.9 | 1372 | 6857 |
| *10AA* datasets (AA) | | | | | | |
| - 1GADBL_100 | 561 | 0.457 | 0.715 | 33.7 | 325 | 490 |
| - RV100_BBA0039 | 807 | 0.416 | 1.000 | 85.3 | 395 | 2696 |
| - RV100_BBA0067 | 410 | 0.783 | 0.922 | 57.5 | 464 | 1092 |
| - RV100_BBA0081 | 353 | 0.863 | 1.000 | 65.2 | 586 | 1682 |
| - RV100_BBA0101 | 509 | 0.784 | 1.000 | 87.9 | 492 | 4081 |
| - RV100_BBA0117 | 460 | 0.754 | 1.000 | 48.4 | 57 | 110 |
| - RV100_BBA0134 | 717 | 0.729 | 1.000 | 85.2 | 470 | 3184 |
| - RV100_BBA0154 | 303 | 0.660 | 0.850 | 59.3 | 518 | 1275 |
| - RV100_BBA0190 | 397 | 0.688 | 1.000 | 64.6 | 886 | 2504 |
| - coli_epi_100 | 320 | 0.583 | 0.871 | 10.7 | 133 | 149 |
| Rec and Res datasets (AA) | | | | | | |
| - Rec | 96,773 | 0.807 | 0.926 | 48.9 | 101 | 205 |
| - Res | 186,802 | 0.735 | 0.908 | 31.2 | 133 | 208 |

#### S3.2 Dataset Generation: INDELible

**Table S2: Indel rate and tree scale parameters for generating the 5000M-het series data, where $l$ is the tree scale parameter, defining the maximum path-length in the non-ultrametric model tree, and $r$ defines the indel rate.**

| Condition | $l$ (tree scale) | $r$ (indel rate) |
|---|---|---|
| 5000M2 | 30 | $4.4 \times 10^{-4}$ |
| 5000M3 | 20 | $3.3 \times 10^{-4}$ |
| 5000M4 | 5 | $1.32 \times 10^{-3}$ |

We present details for generating our new simulated conditions using INDELible [Fletcher and Yang, 2009] below. Our simulation parameters are based on those of the 1000M series of the ROSE simulated dataset [Liu et al., 2009]. We uploaded all of our INDELible control files generating this data to `https://github.com/ThisBioLife/5000M-234-het` to allow easy reproduction.

##### S3.2.1 Model trees

Our model trees were generated by a two-step process. We first generated random 5000-species birth-death trees (one tree per replicate) using the DendroPy [Sukumaran and Holder, 2010] `treesim.birth_death_tree` function, setting the birth rate initially at 1 with the standard deviation of the change to the birth rate set to 0.2 (see the documentation on this Dendropy function on the birth-death process and the implication of this parameter), with a zero death rate. Then, taking only the topology of the tree generated in the prior step, we instructed INDELible to assign branch lengths to the tree such that the resulting tree is both non-ultrametric and has a maximum path-length $l$. We vary $l$ to control the scale of the tree and hence the rate of evolution of the condition. The choice of $l$ across conditions can be seen in Table S2.

##### S3.2.2 Indel length distribution & indel rates

Our model for sequence length heterogeneity assumes a base ("short indel") distribution of indel lengths. In our case, we simply took the "medium" length distribution from the "M"-series ROSE simulated datasets [Liu et al., 2009] as the short indel distribution. Given an indel event, with probability $p = 0.85$, it draws its length from the short indel distribution. Otherwise, it draws its length from a long indel distribution, in our case set to $\mathrm{NB}(130, 0.5)$. The indel length distribution is thus equivalently a mixture distribution of the short indel distribution (prior probability 0.85) and the long indel distribution (prior probability 0.15). We directly computed the probability mass function (PMF) of this mixture distribution, truncated the PMF, and fed the truncated PMF to INDELible as part of the input. The truncated PMF can be found alongside the uploaded control files in the Github repository linked.

Similar to the original ROSE dataset, we also varied the indel rates across conditions, with the choices for the indel rate $r$ shown in Table S2. The indel rates were chosen analogous to the original indel rates of the ROSE dataset.

###### S3.2.3 GTR parameters

**Table S3: GTR matrix parameters for generating the 5000M-het series dataset. The parameters are set according to the parameters for the ROSE simulated data [Liu et al., 2009].**

| GTR Parameters | |
|---|---|
| `T->C` | 1.2619573850882344 |
| `T->A` | 0.14005536945585983 |
| `T->G` | 0.2877830346145434 |
| `C->A` | 0.35766826674033914 |
| `C->G` | 0.3082674310184066 |
| `A->G` | 1 |

The rest of the parameters (e.g., GTR+$\Gamma$ parameters and the initial sequence length) were chosen to be the same as the ROSE simulated dataset generated for the SATé study [Liu et al., 2009] and can be found in the uploaded control files (`https://github.com/ThisBioLife/5000M-234-het`). We also provide the parameters here:

- GTR parameters: see Table S3
- Stationary frequencies (`TCAG`): $0.311475, 0.191363, 0.300414, 0.196748$
- $\alpha$ (for the gamma distribution): 1
- Initial sequence length: 1000

#### S3.3 Dataset Generation: Rec and Res

##### S3.3.1 Background

Software is available that maps mobile DNAs within bacterial and archaeal genomes [Hudson et al., 2015, Mageeney et al., 2020], where each mapping is associated with the sequence of the integrase enzyme that catalyzes the site-specific integration of the mobile DNA.

##### S3.3.2 Generation

Protein sequences were taken from 350,378 bacterial and archaeal genome sequences (see `https://databank.illinois.edu/datasets/IDB-2419626`) using Prodigal [Hyatt et al., 2010]. Serine recombinases were identified using the Pfam HMMs [Mistry et al., 2021] Resolvase (Res) for the catalytic domain and Recombinase (Rec) for the integrase-specific domain. Seed sequences are included for evaluation later (112 and 66 Pfam seed sequences for Res and Rec, respectively).

Link to the Res seed sequences: `http://pfam.xfam.org/family/PF00239#tabview=tab3`

Link to the Rec seed sequences: `http://pfam.xfam.org/family/PF07508#tabview=tab3`

**Sequence trimming** Two separate sequence datasets (i.e., Rec and Res) were prepared for the serine recombinases trimmed either according to the Recombinase or Resolvase HMM hits.

##### S3.3.3 Evaluation

We compare the estimated alignments to the seed Pfam alignments for both Rec and Res to obtain SPFN, SPFP, and the expansion score (see main text for definitions).

### S4 Computational issues for MAFFT-linsi-add and MAFFT-add

#### S4.1 Experiment 0: MAFFT-linsi-add scalability issue on 5000-taxon datasets

In Experiment 0, we limited the runtime to 12 hours, 64 GB of memory, and 16 cores. We varied the number of queries to be added in this experiment. However, when we tried running MAFFT-linsi --add (MAFFT-linsi-add) on our training 5000M2 datasets with the full set of 4000 query sequences to a 1000-taxon full-length backbone alignment, we encountered either out-of-memory issues (64 GB memory limit) or crashes.

The out-of-memory error message looks like the following:

```
slurmstepd: error: Detected 1 oom-kill event(s) in
StepId=5376434.batch cgroup. Some of your processes may have been
killed by the cgroup out-of-memory handler.
```

The crash error message looks like the following:

```
Command exited with non-zero status 1
```

#### S4.2 Experiment 2: MAFFT-add out-of-memory issues

In Experiment 2, we allowed 24 hours and 128 GB memory, as well as 16 cores. We ran MAFFT-add (default option) for all three ways of defining the backbone sequences on the Res dataset with $\sim 186k$ sequences. This analysis produced out-of-memory issues for MAFFT--add, with the following error message:

```
slurmstepd: error: *** JOB 8737367 ON ccc0281 CANCELLED AT
    2023-05-05T17:47:50 ***
slurmstepd: error: Detected 1 oom-kill event(s) in StepId=8737367.
    batch. Some of your processes may have been killed by the
    cgroup out-of-memory handler.
```

### S5 Additional Figures

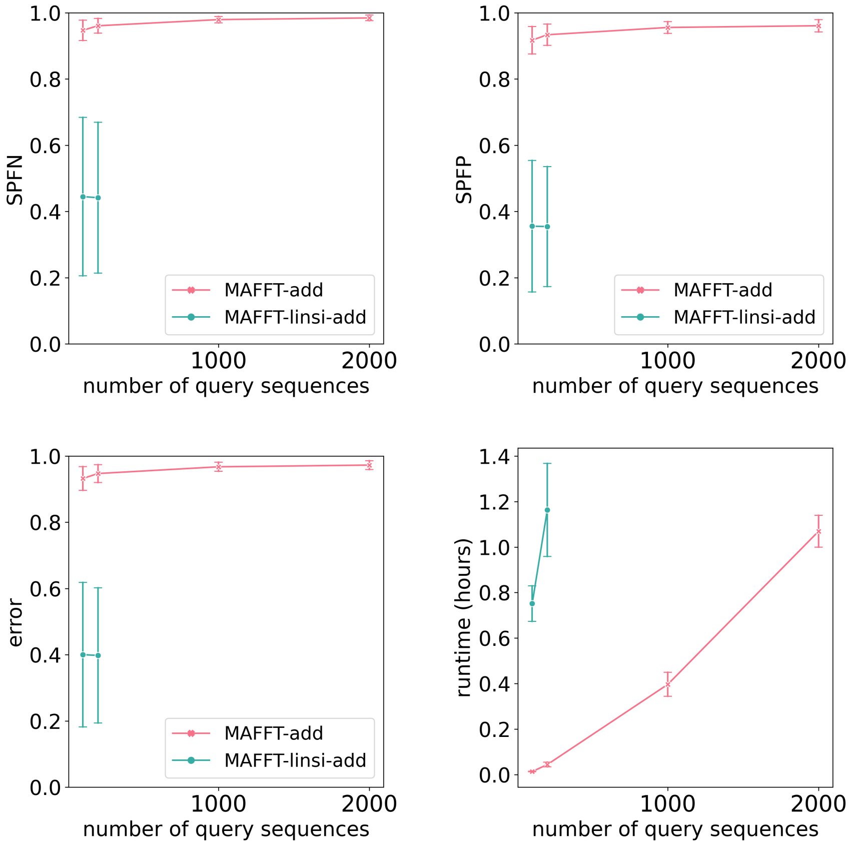

**Figure S1: Experiment 0: SPFN (top left), SPFP (top right), alignment error (average of SPFN and SPFP, bottom left), and runtime in hours (bottom right) of MAFFT-add and MAFFT-linsi-add for adding 100, 200, 1000, or 2000 sequences to a 1000-taxon backbone alignment. The dataset used is 5000M2-het with 10 replicates, where 1000 full-length sequences are randomly selected and aligned with MAGUS [Smirnov and Warnow, 2021] to form the backbone alignment. Averages over ten replicates are shown. Error bars shown for alignment errors are standard errors and standard deviations for runtime. We exclude replicate 4 because MAFFT-linsi-add encountered out-of-memory issues when adding 100 or 200 query sequences. Additionally, MAFFT-linsi-add either encountered out-of-memory issues or did not complete within 12 hours when adding 1000 or 2000 query sequences and thus is not shown.**

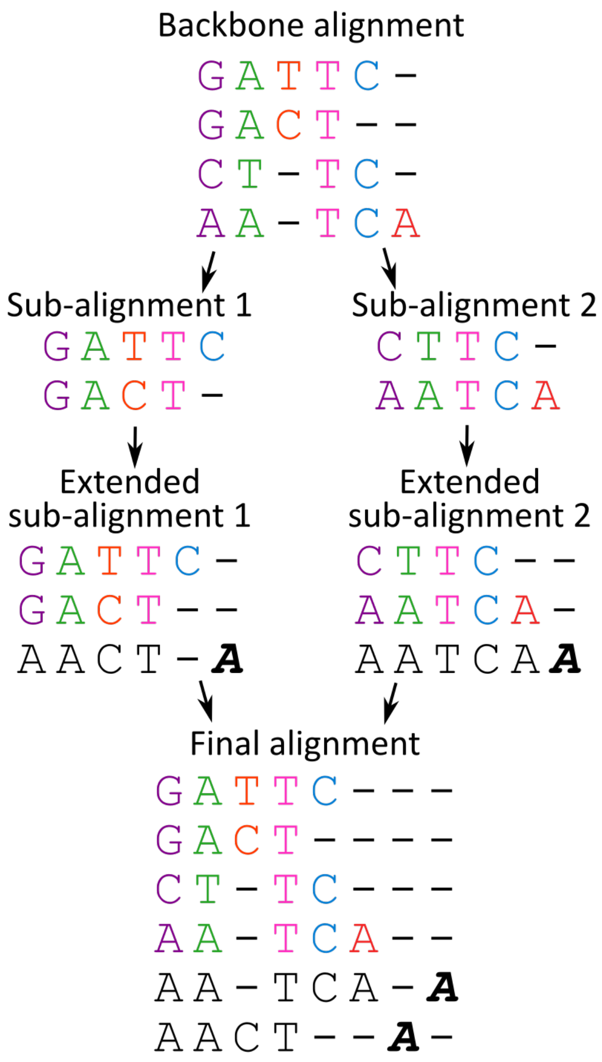

**Figure S2: A simple example of how transitivity merging works when merging two extended subalignments, each adding one query new sequence. Insertion letters are marked in bold and italicized. The two insertion letters, marked in the final columns of the two extended subalignments, are put into separate columns in the final alignment. See text for additional discussion.**

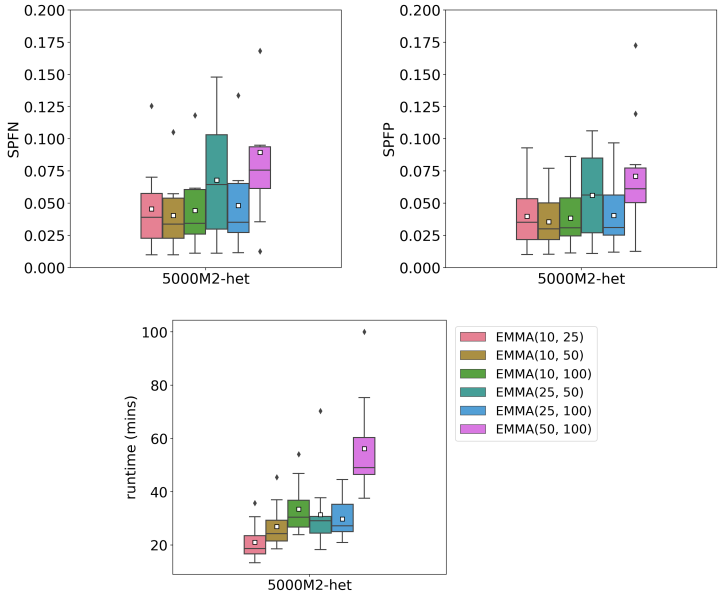

**Figure S3: Experiment 1: We show alignment error SPFN (top left) and SPFP (top right) as well as runtime (bottom) for EMMA run with different settings of $l$ and $u$ (the algorithmic parameters governing subset size). Parameters $l$ and $u$ are drawn from $\{10, 25, 50, 100\}$, and with $l < u$. This experiment was performed with a backbone of 1000 randomly selected sequences of 10 replicates of the 5000M2-het model condition. Alignment error is calculated on query sequences only, and white squares mark the averages.**

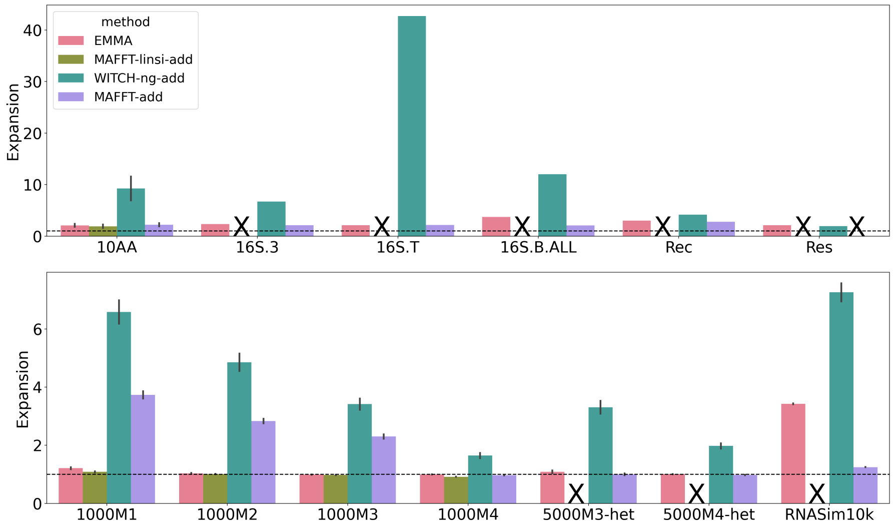

**(a) Expansion score**

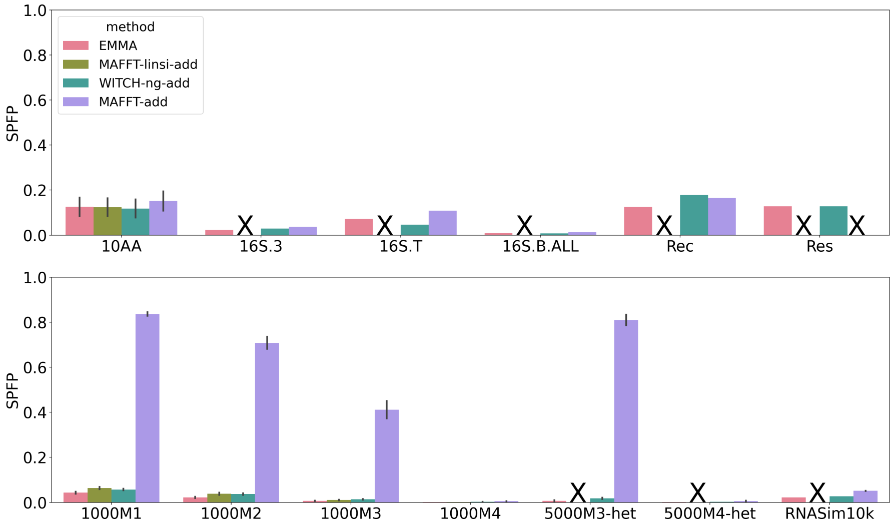

**(b) SPFP**

**Figure S4: Experiment 2: Expansion score (top) and SPFP (bottom) when adding to large random backbone alignments. In each sub-figure, the top panel denotes biological datasets, and the bottom panel denotes simulated datasets. The horizontal dashed line indicates an expansion score of 1. MAFFT-linsi-add failed to finish within 24 hours for datasets except for 10AA and ROSE 1000M1-4 and was not run on Rec and Res due to their large numbers of sequences. MAFFT-add encountered out-of-memory issues on the Res dataset, as discussed in Section S4. Failed runs, or runs not attempted, are marked with "X".**

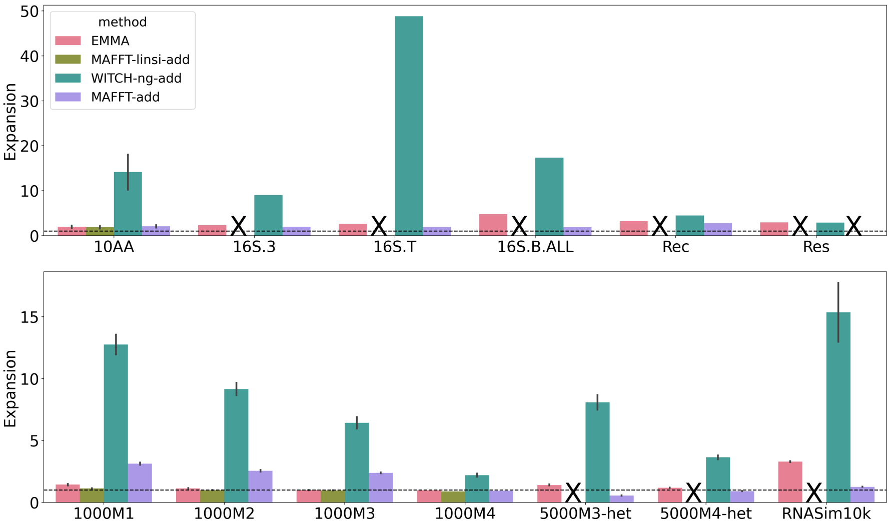

**(a) Expansion score**

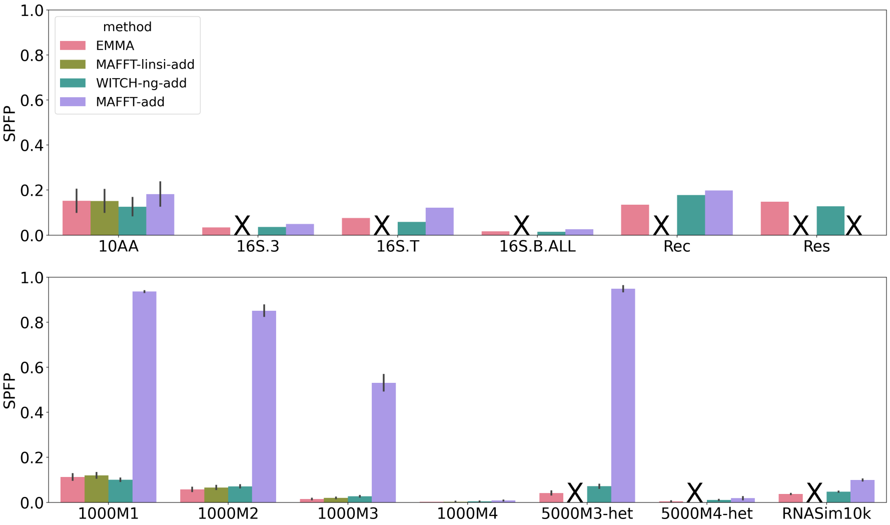

**(b) SPFP**

**Figure S5: Experiment 2: Expansion score (top) and SPFP (bottom) when adding to small random backbone alignments. In each sub-figure, the top panel denotes biological datasets, and the bottom panel denotes simulated datasets. The horizontal dashed line indicates an expansion score of 1. MAFFT-linsi-add failed to finish within 24 hours for datasets except for 10AA and ROSE 1000M1-4 and was not run on Rec and Res due to their large numbers of sequences. MAFFT-add encountered out-of-memory issues on the Res dataset (Section S4). Failed runs, or runs not attempted, are marked with "X".**

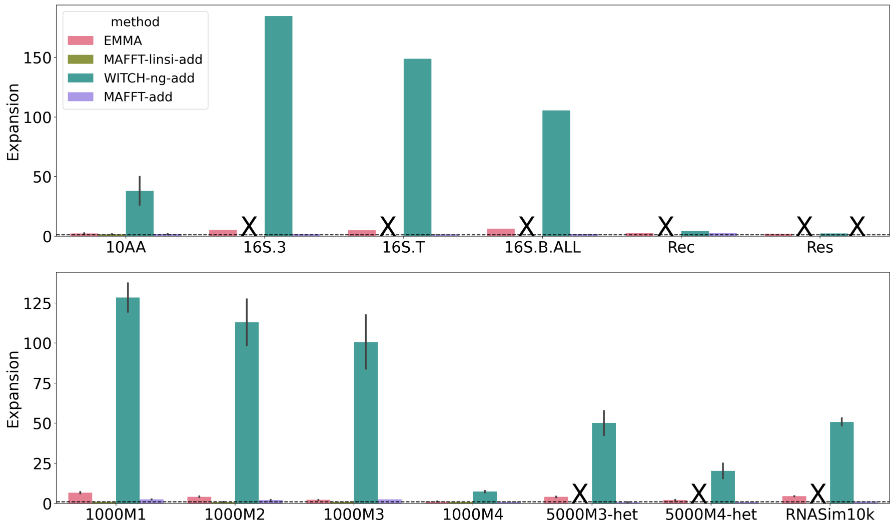

**(a) Expansion score**

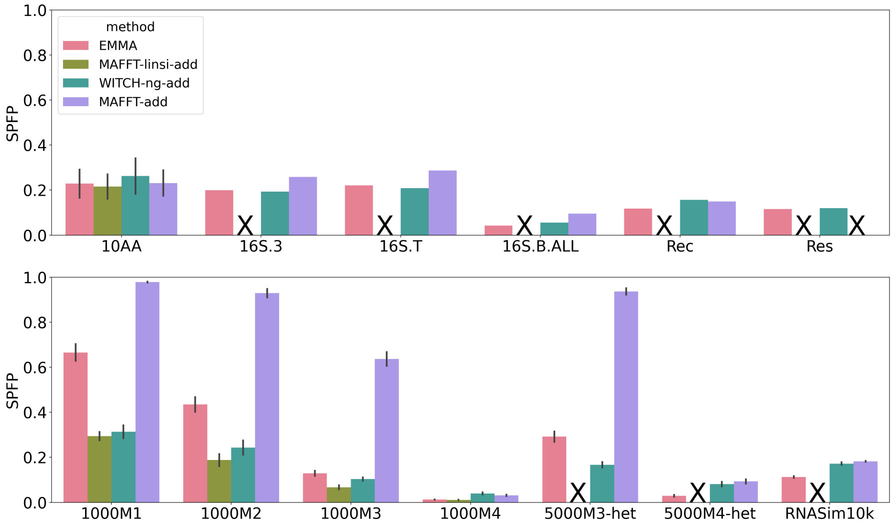

**(b) SPFP**

**Figure S6: Experiment 2: Expansion score (top) and SPFP (bottom) when adding to clade-based backbone alignments. In each sub-figure, the top panel denotes biological datasets, and the bottom panel denotes simulated datasets. The horizontal dashed line indicates an expansion score of 1. MAFFT-linsi-add failed to finish within 24 hours for datasets except for 10AA and ROSE 1000M1-4 and was not run on Rec and Res due to their large numbers of sequences. MAFFT-add encountered out-of-memory issues on the Res dataset (Section S4). Failed runs, or runs not attempted, are marked with "X".**

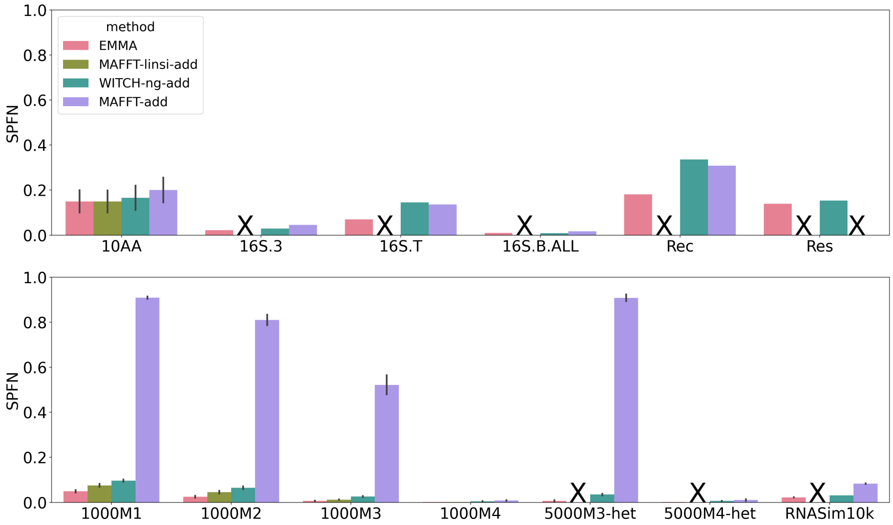

**Figure S7: Experiment 2: SPFN when adding to large random backbone alignments. The top panel denotes biological datasets, and the bottom panel denotes simulated datasets. MAFFT-linsi-add failed to finish within 24 hours for datasets except for 10AA and ROSE 1000M1-4 and was not run on Rec and Res due to their large numbers of sequences. MAFFT-add encountered out-of-memory issues on the Res dataset (Section S4). Failed runs, or runs not attempted, are marked with "X".**

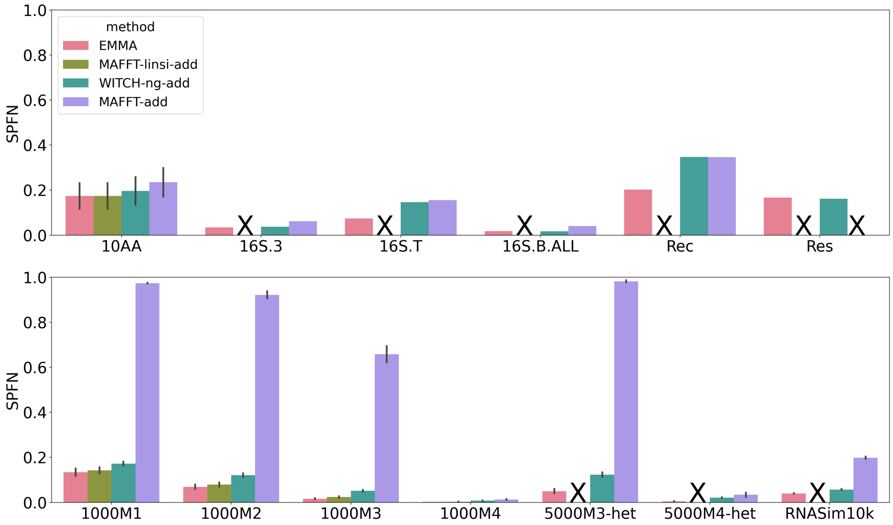

**Figure S8: Experiment 2: SPFN when adding to small random backbone alignments. The top panel denotes biological datasets, and the bottom panel denotes simulated datasets. MAFFT-linsi-add failed to finish within 24 hours for datasets except for 10AA and ROSE 1000M1-4 and was not run on Rec and Res due to their large numbers of sequences. MAFFT-add encountered out-of-memory issues on the Res dataset (Section S4). Failed runs, or runs not attempted, are marked with "X".**

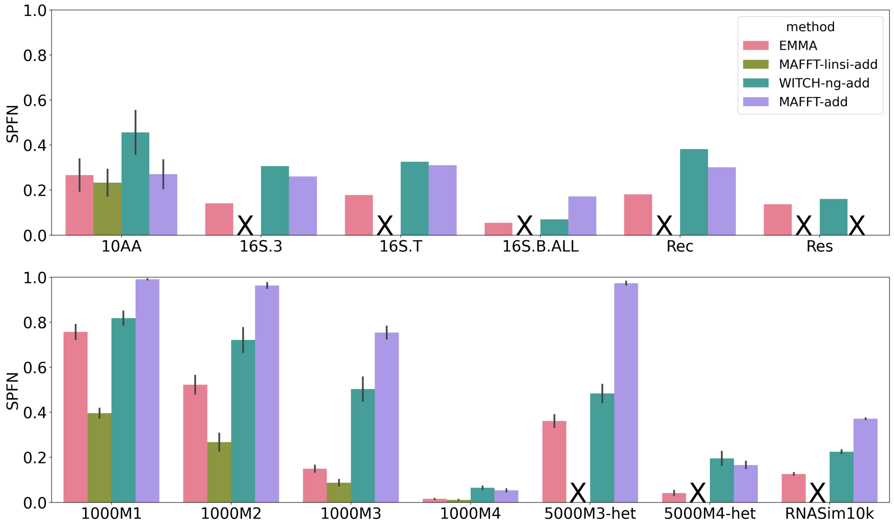

**Figure S9: Experiment 2: SPFN when adding to clade-based backbone alignments. The top panel denotes biological datasets, and the bottom panel denotes simulated datasets. MAFFT-linsi-add failed to finish within 24 hours for datasets except for 10AA and ROSE 1000M1-4 and was not run on Rec and Res due to their large numbers of sequences. MAFFT-add encountered out-of-memory issues on the Res dataset (Section S4). Failed runs, or runs not attempted, are marked with "X".**

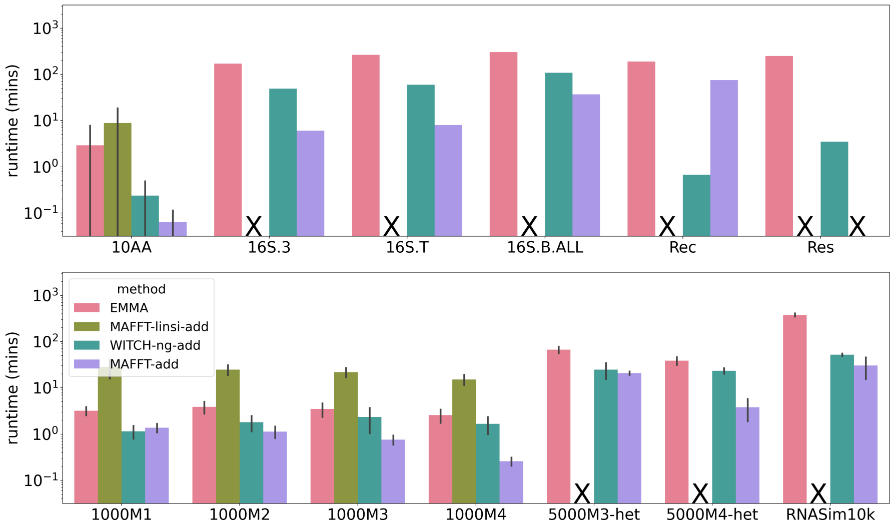

**Figure S10: Experiment 2: Runtime (log-scale) in minutes when adding to small random backbone alignments. The top panel denotes biological datasets, and the bottom panel denotes simulated datasets. MAFFT-linsi-add failed to finish within 24 hours for datasets except for 10AA and ROSE 1000M1-4 and was not run on Rec and Res due to their large numbers of sequences. MAFFT-add encountered out-of-memory issues on the Res dataset (Section S4). Failed runs, or runs not attempted, are marked with "X". Error bars indicate standard deviation.**

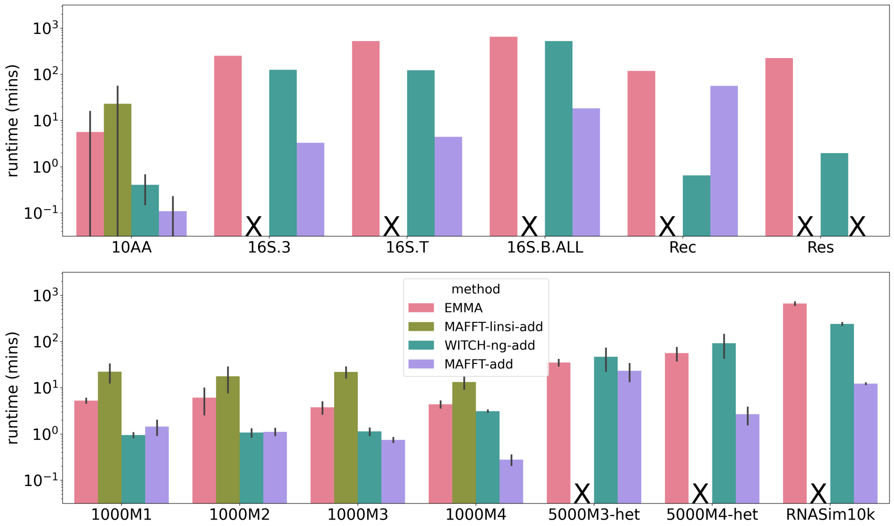

**Figure S11: Experiment 2: Runtime (log-scale) in minutes when adding to clade-based backbone alignments. The top panel denotes biological datasets, and the bottom panel denotes simulated datasets. MAFFT-linsi-add failed to finish within 24 hours for datasets except for 10AA and ROSE 1000M1-4 and was not run on Rec and Res due to their large numbers of sequences. MAFFT-add encountered out-of-memory issues on the Res dataset (Section S4). Failed runs, or runs not attempted, are marked with "X". Error bars indicate standard deviation.**

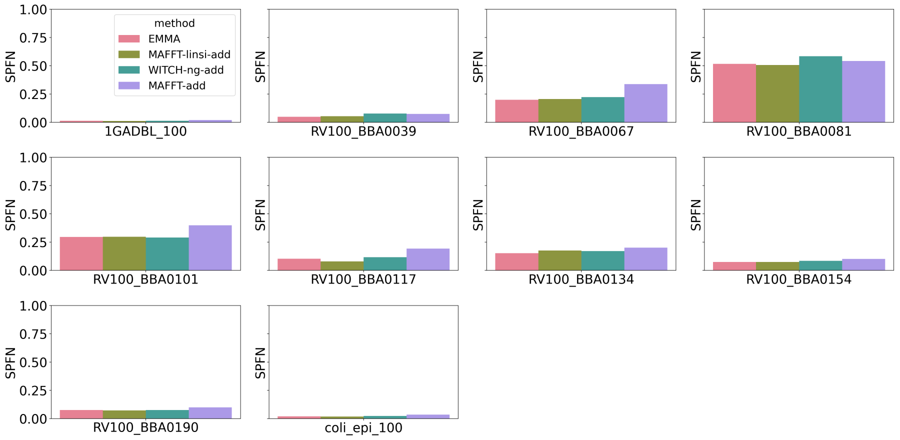

**Figure S12: Experiment 2: Individual SPFN for 10AA datasets when adding to large random backbone alignments.**

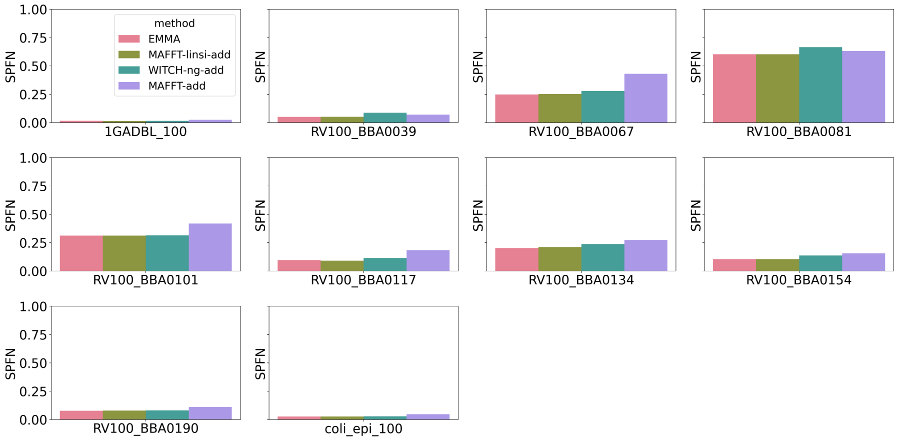

**Figure S13: Experiment 2: Individual SPFN for 10AA datasets when adding to small random backbone alignments.**

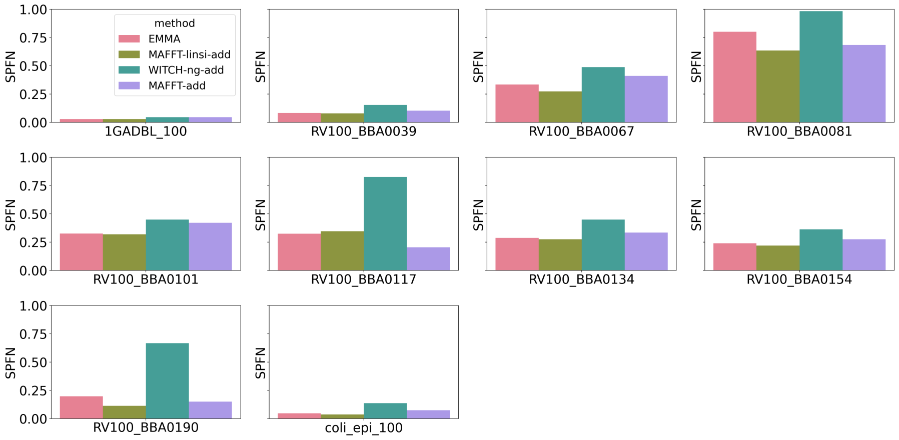

**Figure S14: Experiment 2: Individual SPFN for 10AA datasets when adding to large clade-based alignments.**

#### S6 Additional Tables

**Table S4: Average expansion scores of EMMA, WITCH-ng-add, MAFFT --add, and MAFFT-linsi --add for different model conditions on all datasets. "X" denotes that the method failed due to runtime (24-hour limit) or out-of-memory issues (128 GB). All values are rounded to the nearest tenth.**

| | EMMA | WITCH-ng-add | MAFFT --add | MAFFT-linsi --add |
|---|---|---|---|---|
| **large random backbone** | | | | |
| 10AA | 2.0 | 9.3 | 2.2 | 2.1 |
| 16S.3 | 2.1 | 6.7 | 2.1 | X |
| 16S.T | 2.1 | 42.7 | 2.2 | X |
| 16S.B.ALL | 3.9 | 12.0 | 2.1 | X |
| Rec | 3.3 | 4.2 | 2.8 | X |
| Res | 1.7 | 2.0 | X | X |
| 1000M1 | 1.2 | 6.6 | 3.7 | 1.1 |
| 1000M2 | 1.0 | 4.9 | 2.8 | 1.0 |
| 1000M3 | 1.0 | 3.4 | 2.3 | 1.0 |
| 1000M4 | 1.0 | 1.6 | 1.0 | 0.9 |
| 5000M3-het | 1.0 | 3.3 | 1.0 | X |
| 5000M4-het | 1.0 | 2.0 | 1.0 | X |
| RNASim10k | 2.8 | 7.3 | 1.2 | X |
| **small random backbone** | | | | |
| 10AA | 2.0 | 14.1 | 2.1 | 1.9 |
| 16S.3 | 3.0 | 9.1 | 2.0 | X |
| 16S.T | 3.1 | 48.9 | 1.9 | X |
| 16S.B.ALL | 5.6 | 17.4 | 1.9 | X |
| Rec | 3.2 | 4.5 | 2.8 | X |
| Res | 2.9 | 2.9 | X | X |
| 1000M1 | 1.6 | 12.8 | 3.1 | 1.1 |
| 1000M2 | 1.3 | 9.2 | 2.5 | 1.0 |
| 1000M3 | 1.1 | 6.4 | 2.4 | 1.0 |
| 1000M4 | 1.0 | 2.2 | 0.9 | 0.9 |
| 5000M3-het | 2.0 | 8.1 | 0.5 | X |
| 5000M4-het | 1.6 | 3.6 | 0.9 | X |
| RNASim10k | 3.2 | 15.4 | 1.3 | X |
| **large clade-based backbone** | | | | |
| 10AA | 1.7 | 38.1 | 1.7 | 1.5 |
| 16S.3 | 4.0 | 184.7 | 1.6 | X |
| 16S.T | 4.8 | 148.8 | 1.5 | X |
| 16S.B.ALL | 6.3 | 105.6 | 1.7 | X |
| Rec | 2.3 | 4.3 | 2.7 | X |
| Res | 1.9 | 2.2 | X | X |
| 1000M1 | 2.0 | 128.4 | 2.5 | 0.9 |
| 1000M2 | 1.5 | 112.9 | 2.1 | 0.9 |
| 1000M3 | 1.3 | 100.7 | 2.5 | 0.9 |
| 1000M4 | 1.1 | 7.3 | 0.9 | 0.9 |
| 5000M3-het | 2.2 | 50.1 | 0.7 | X |
| 5000M4-het | 2.0 | 20.3 | 0.9 | X |
| RNASim10k | 3.9 | 50.8 | 1.2 | X |

**Table S5: Memory usage (mean, minimum, and maximum in GB) of EMMA, WITCH-ng-add, and MAFFT--add for different model conditions on all datasets. For min/max memory usage, we also show the specific dataset in parentheses. The maximum memory usage of MAFFT--add is the memory limit (128 GB) for Res, and led to out-of-memory crashes. MAFFT-linsi--add only ran on the datasets with at most 1000 sequences, and so its memory usage is not comparable to the other methods. All values are rounded to the nearest hundreds.**

| backbone type | method | memory mean | memory min | memory max |
|---|---|---|---|---|
| large random | EMMA | 0.57 | 0.04 (RV100_BBA0117) | 4.93 (16S.3) |
| large random | WITCH-ng-add | 0.93 | 0.01 (RV100_BBA0117) | 8.06 (16S.B.ALL) |
| large random | MAFFT--add | 2.53 | 0.03 (coli_epi_100) | 127.95 (Res) |
| small random | EMMA | 1.21 | 0.03 (RV100_BBA0117) | 6.16 (RV100_BBA0101) |
| small random | WITCH-ng-add | 1.23 | 0.01 (RV100_BBA0117) | 11.23 (16S.B.ALL) |
| small random | MAFFT--add | 2.52 | 0.03 (coli_epi_100) | 127.95 (Res) |
| clade-based | EMMA | 1.23 | 0.04 (coli_epi_100) | 4.47 (16S.3) |
| clade-based | WITCH-ng-add | 0.92 | 0.00 (RV100_BBA0117) | 12.84 (16S.B.ALL) |
| clade-based | MAFFT--add | 2.66 | 0.03 (coli_epi_100) | 127.95 (Res) |
